# Glucose Starvation Sensing through Membrane Remodeling at the Nucleus-Vacuole Junction Coordinates Ergosterol Biosynthesis

**DOI:** 10.1101/2025.05.29.656913

**Authors:** Shintaro Fujimoto, Yasushi Tamura

## Abstract

Membrane contact sites (MCSs), where organelle membranes come into close proximity, function as dynamic hubs for lipid metabolism in response to metabolic and stress signals. In yeast, the nucleus-vacuole junction (NVJ) expands during glucose starvation through the recruitment of stress-specific proteins, but the underlying mechanisms and physiological significance remained unclear. Here, we identify the yeast INSIG homologs Nsg1, Nsg2, along with the aspartyl protease Ypf1 as glucose starvation-specific NVJ-localized factors. Ypf1 promotes the recruitment of INSIG proteins and the HMG-CoA reductases Hmg1 and Hmg2 to the NVJ, likely in response to nuclear membrane remodeling induced by changes in very long-chain fatty acid (VLCFA) metabolism. We further demonstrate that this membrane remodeling destabilizes Nsg1, promoting the NVJ localization and activation of Hmg1, while stabilizing Nsg2, which counteracts Hmg1 activation. These findings reveal that Nsg1 and Nsg2, previously known as regulators of Hmg2 stability, also act as negative regulators of Hmg1. Based on these results, we propose a model in which VLCFAs modulate the physical properties of the nuclear membrane, enabling cells to sense glucose starvation and regulate sterol metabolism during glucose starvation. Overall, our study provides new insights into MCS-mediated metabolic stress responses and highlights the role of membrane properties as active regulators of lipid homeostasis.

## Introduction

Membrane contact sites (MCSs), where different biological membranes are in close proximity, have garnered significant attention as critical hubs for lipid and ion transport between adjacent organelles (*1–7*). Recent studies have revealed that MCSs play important roles in metabolic regulation and cellular stress responses through dynamic changes in their number and size. For instance, the ERMES (ER-Mitochondria Encounter Structure) complex, which forms MCSs between the ER and mitochondria in yeast (*8*), undergoes structural remodeling in response to nutrient availability and ER stress (*9–11*). Under ER stress conditions, ERMES foci, clusters of oligomerized ERMES complexes, dissociate, thereby suppressing phospholipid transport from the ER to mitochondria. This suppression of phospholipid transport appears to contribute to ER membrane expansion, alleviating ER stress (*10*). Furthermore, the number of ERMES foci and vCLAMPs, the mitochondria-vacuole MCS, is reciprocally regulated in response to carbon source availability and the activity of either ERMES or vCLAMP (*9*, *12*, *13*). The absence of vCLAMPs leads to an increase in ERMES foci, whereas ERMES deficiency promotes vCLAMP formation. Additionally, under respiratory growth conditions, vCLAMP formation is reduced, while ERMES foci increase, suggesting that these MCSs are dynamically regulated in response to metabolic signals.

The nucleus-vacuole junction (NVJ), an MCS between the nuclear and vacuolar membranes, is also highly responsive to nutritional conditions (*14–17*). The NVJ is formed through the direct interaction of the nuclear membrane protein Nvj1 and the vacuolar membrane protein Vac8 (*18*, *19*). Several proteins, including Osh1, Tsc13, and Nvj2, accumulate at the NVJ in an Nvj1-dependent manner (*20–22*). Given that Osh1 and Nvj2 contain lipid transport-related domains such as an oxysterol-binding domain and an SMP domain, respectively and that Tsc13 is involved in the elongation of very long-chain fatty acids (VLCFAs) (*21–23*), the NVJ serves as a crucial site for lipid metabolism. In addition, the NVJ facilitates piecemeal microautophagy of the nucleus (PMN), a type of selective autophagy in which portions of the nucleus are directly engulfed by the vacuole upon nitrogen starvation or rapamycin treatment (*24–26*). Moreover, Mdm1 and its paralog Nvj3 localize to the NVJ in an Nvj1-independent manner (*27*). Mdm1 recruits Faa1, a fatty acid-metabolizing enzyme, to the NVJ and is involved in lipid droplet synthesis (*28*).

The NVJ also plays a critical role in the cellular response to glucose starvation (GS). During GS, Snd3, a factor involved in ER membrane protein transport (*29*), accumulates at the NVJ, contributing to NVJ expansion (*30*). Additionally, Hmg1, the rate-limiting enzyme in sterol biosynthesis, accumulates at the NVJ in a GS-dependent manner, where it oligomerizes to enhance ergosterol synthesis (*31*). However, the mechanisms by which these factors sense GS and relocate from the ER or nuclear membrane to the NVJ remain largely unknown. Furthermore, the physiological significance of the NVJ remodeling under GS conditions remains to be elucidated.

Here, we report that the yeast homologs of mammalian INSIG (Insulin Induced Gene) proteins, Nsg1 and Nsg2 (*32*), along with the intramembrane aspartyl protease Ypf1 (*33*), accumulate at the NVJ in a GS-dependent manner and play a crucial role in regulating ergosterol synthesis. Ypf1 appears to mediate the recruitment of Nsg1, Nsg2, Hmg1, and Hmg2 to the NVJ through its GS-dependent structural changes. We further demonstrate that GS leads to the destabilization of Nsg1 and the stabilization of Nsg2. Importantly, our results reveal that both Nsg1 and Nsg2 function as negative regulators of Hmg1, and that their opposing stabilities under GS conditions enable coordinated regulation of Hmg1 activity to maintain proper ergosterol levels. Consistently, simultaneous deletion of Nsg1 and Nsg2 leads to a dramatic accumulation of ergosterol esters and squalene, indicating that the loss of both negative regulators results in hyperactivation of Hmg1 and excessive sterol biosynthesis. Notably, we also show that these NVJ remodeling events are driven by GS-dependent changes in VLCFA metabolism. Together, these findings provide novel insights into the molecular mechanisms underlying metabolic adaptation to GS, mediated by the dynamic NVJ reorganization, and also highlight the potential of this regulatory system for applications in yeast-based production of commercially valuable lipids.

## Results

### Searching for Novel NVJ-Localized Factors Using CsFiND

We previously developed CsFiND (<u>C</u>omplementation a<u>s</u>say using fusion of split-GFP a<u>n</u>d TurboI<u>D</u>), a proximity labeling technique that specifically targets MCSs (*34*). In this system, fusion proteins composed of tandemly linked split fragments of GFP and TurboID are expressed on different organelle membranes. When the two membranes come into close proximity at MCSs, complete GFP and TurboID are reconstituted, enabling both visualization of the MCS and biotinylation of MCS-localized proteins (Fig. 1A). To identify novel proteins localized at the NVJ, we adapted the CsFiND system to this MCS. Specifically, we constructed two CsFiND proteins: one consisting of full-length Dpp1, the N-terminal fragment of TurboID, a V5 tag, and the C-terminal fragment of GFP (GFP11); and the other consisting of full-length Ifa38, the N-terminal fragment of GFP (GFP1–10), a FLAG tag, and the C-terminal fragment of TurboID (Fig. 1B). Dpp1 and Ifa38 have been successfully used in our previous studies to localize split-GFP fragments to the ER and vacuolar membranes for visualizing the MCSs (*35*, *36*). Confocal microscopy revealed that the reconstituted GFP signals generated by CsFiND were colocalized with Nvj1-mCherry, a known NVJ marker, indicating that the CsFiND components were successfully reassembled specifically at the NVJ (Fig. 1C).

**Fig. 1.**
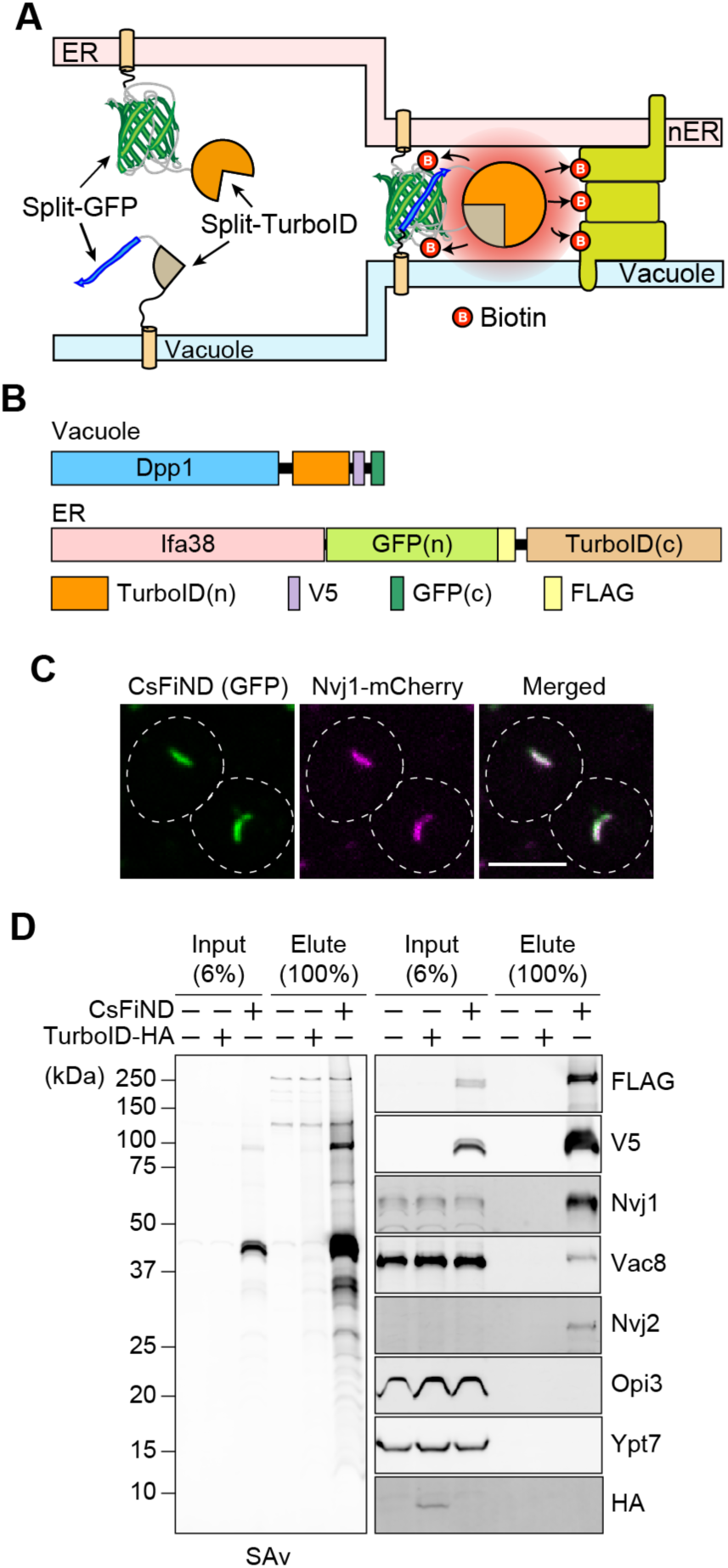
CsFiND for identification of NVJ-localized proteins. (**A**) A schematic diagram illustratinghow CsFiND proteins biotinylate NVJ-localized proteins. (**B**) Schematic representation of the CsFiND constructs used in this study. (**C**) Yeast cells expressing CsFiND proteins and Nvj1-mCherry were observed by confocal fluorescence microscopy. Scale Bar, 5 µm. (**D**) The biotinylated proteins were purified from membrane fractions isolated from wild-type cells with or without the expression of CsFiND proteins or TurboID-HA and subjected to streptavidin blotting and immunoblotting using the indicated antibodies.

Next, we examined whether NVJ-localized proteins were biotinylated by CsFiND. Wild-type cells were cultured in the presence of biotin, with or without expression of either CsFiND or cytosolic TurboID-HA. Heavy membrane fractions were isolated to remove soluble endogenous biotinylated proteins, which could interfere with subsequent mass spectrometry-based protein identification (Fig. 1D and S1A). The membrane fractions were solubilized and subjected to streptavidin-based purification. Purified proteins were detected using streptavidin-Cy5 and antibodies against known NVJ proteins (Fig. 1D). We found that biotinylated proteins were barely detectable in membrane fractions from cells lacking CsFiND or expressing TurboID-HA alone (Fig. 1D). In contrast, membranes from cells expressing CsFiND showed numerous biotinylated proteins visualized by streptavidin-Cy5. Notably, CsFiND proteins containing FLAG and V5 tags, as well as known NVJ-localized proteins such as Nvj1, Vac8, and Nvj2, were enriched in streptavidin-purified samples, whereas ER-localized Opi3 and the vacuolar membrane protein Ypt7 were not (Fig. 1D). These results indicate that CsFiND selectively biotinylates proteins localized at the NVJ.

We then subjected the purified biotinylated proteins to LC-MS/MS analysis, which identified several previously reported NVJ-localized proteins, including Nvj1, Nvj2, Vac8, Tsc13, Osh1, Mdm1, Lam6, Hmg1, Hmg2, and Faa1 (Fig. S1B). We examined the previously reported subcellular localizations of the identified proteins using databases such as YeastRGB (*37*) and LoQAtE (*38*), and selected proteins showing even faint NVJ-like localization for further analysis. These included Nsg1 and Nsg2, the yeast homologs of mammalian INSIG (Insulin Induced Gene) proteins (*32*), as well as Ypf1, an asparagine- processing peptidase (*33*).

### Nsg1, Nsg2, and Ypf1 localize to NVJ during glucose starvation

To analyze subcellular localizations of Ypf1, Nsg1 and Nsg2, we chromosomally expressed Ypf1-GFP, mCherry-Nsg1, or GFP-Nsg2 together with GFP or mCherry-fused Nvj1 as an NVJ marker. Our microscopy analyses revealed that Ypf1, Nsg1, and Nsg2 were primarily localized to the nuclear membrane under nutrient-rich conditions, and did not accumulate at the NVJ labeled with Nvj1 (Fig. 2A, Log). Under a nitrogen starvation condition, Ypf1, Nsg1, and Nsg2 did not show NVJ localization while the NVJ region labeled with Nvj1 expanded (Fig. 2A, NS). Interestingly, Ypf1, Nsg1, and Nsg2 clearly accumulated at the expanded NVJ during GS. These results indicate that Ypf1, Nsg1, and Nsg2 are GS-specific NVJ factors (Fig. 2A, GS).

**Fig. 2.**
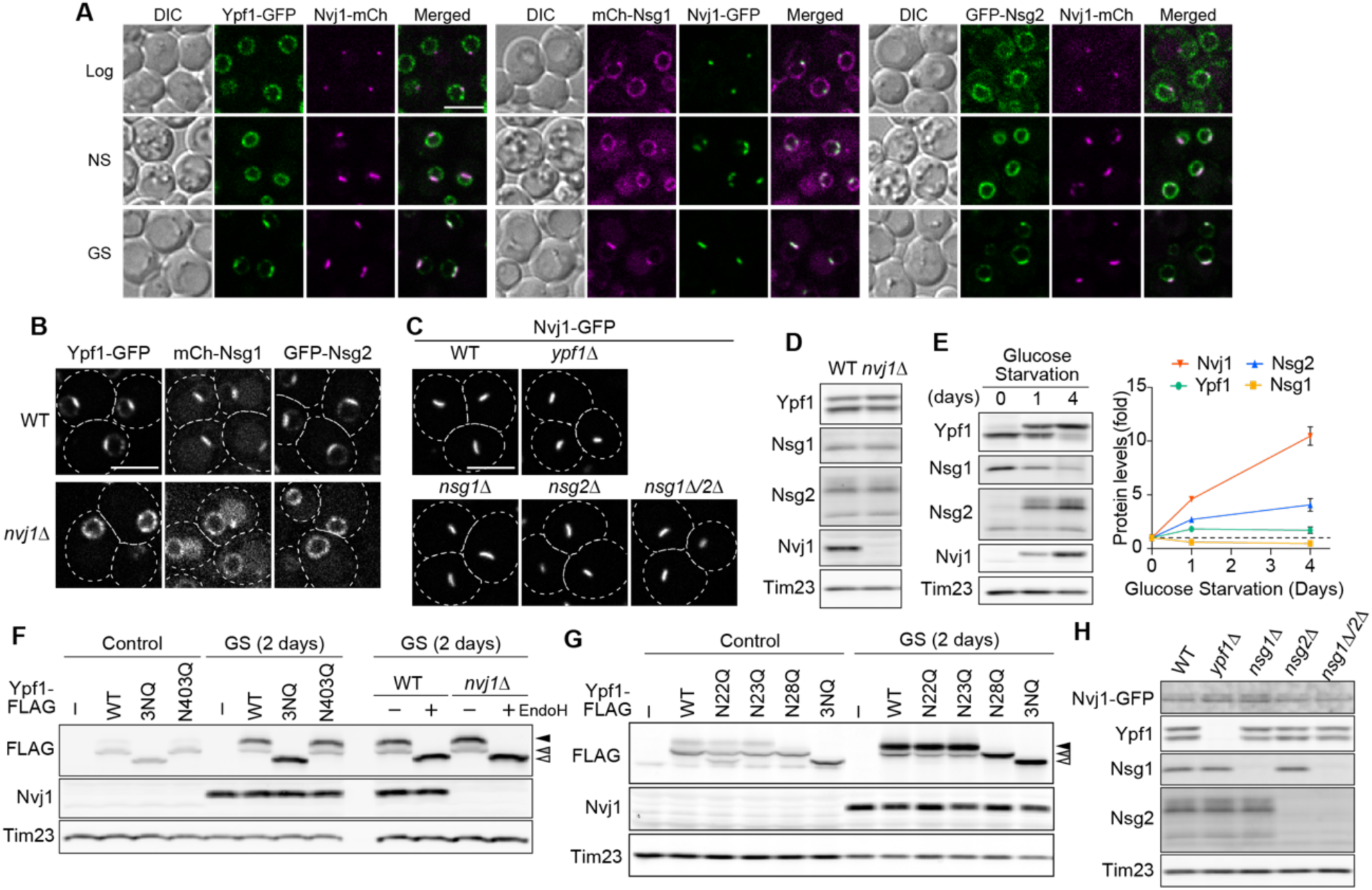
Ypf1, Nsg1, and Nsg2 accumulate at the NVJ under GS conditions. (**A**) Yeast cells co-expressing Ypf1-GFP/Nvj1-mCherry, mCherry-Nsg1/Nvj1-GFP, or GFP-Nsg2/Nvj1-mCherry were observed by confocal microscopy under the indicated culture conditions. Log, NS, and GS refer to cells grown in logarithmic phase in YPD, or subjected to nitrogen or glucose starvation for 24 hours, respectively. (**B**) Wild-type and nvj1Δ cells expressing Ypf1-GFP, mCherry-Nsg1, or GFP-Nsg2 were imaged by confocal fluorescence microscopy after a 24-hour incubation in GS medium. (**C**) The indicated yeast cells expressing Nvj1-GFP were observed by confocal fluorescence microscopy after a 24-hour incubation in GS medium. (**D**) Total cell lysates from wild-type and *nvj1*Δ cells after a 24-hour incubation in GS medium were analyzed by immunoblotting using the indicated antibodies. (**E**) Total cell lysates were prepared from wild-type cells grown in GS medium for the indicated periods of time and analyzed by immunoblotting. Line graph shows protein levels relative to those in cells without GS treatment. Values are means ± S.E. (n = 3). (**F, G**) Glycosylation patterns of Ypf1-FLAG and its mutants were analyzed by immunoblotting using whole cell lysates prepared from cells with or without a 2-day GS treatment. (**H**) Total cell lysates from the cells shown in (**C**) were analyzed by immunoblotting. All images shown in (**A**), (**B**), and (**C**) are single focal plane. Scale bar, 5 μm.

How do these factors accumulate at the NVJ in a GS-dependent manner? It has been reported that Nvj1 is critical for the NVJ partitioning of Hmg1, which is known as a GS-specific NVJ factor(*31*). We thus analyzed the localizations of Ypf1, Nsg1, and Nsg2 in *nvj1Δ* cells. Similar to the case of Hmg1, the loss of Nvj1 abolished the NVJ partitioning of Nsg1, Nsg2 and Ypf1, indicating that the GS-induced NVJ localization of Ypf1, Nsg1, and Nsg2 depends on Nvj1 (Fig. 2B). In contrast, loss of Ypf1, Nsg1, or Nsg2 did not affect the NVJ localization of Nvj1 (Fig. 2C).

We next examined whether the loss of Nvj1 affected the steady state levels of Ypf1, Nsg1, and Nsg2. To this end, we performed immunoblotting of total cell lysates prepared from wild-type and *nvj1Δ* cells grown in GS conditions. The immunoblotting showed that the loss of Nvj1 did not affect the expression levels of Ypf1, Nsg1, and Nsg2 (Fig. 2D). However, we noticed that GS greatly changed the steady state levels and apparent molecular weights of Nvj1, Nsg1, Nsg2, and Ypf1 (Fig. 2E). Specifically, during GS, the level of Nsg1 decreased while those of Nvj1 and Nsg2 increased, indicating contrasting stability regulation. Moreover, GS led to an increase in the apparent molecular weight of Nvj1, Nsg2, and Ypf1, suggesting that they undergo GS-dependent modifications. Previous phosphoproteome studies indicated that Nvj1 and Nsg2 were phosphorylated(*39–41*). Phos-tag gel electrophoresis revealed that the apparent molecular weight of Nvj1 increased markedly during GS, and this shift was reversed to the pre-stress mobility upon phosphatase treatment (Fig. S2A). Similarly, a higher molecular weight band of Nsg2 observed during GS was no longer detectable after phosphatase treatment (Fig. S2A). These results indicate that both Nvj1 and Nsg2 are specifically phosphorylated in response to GS. Since the apparent size of Ypf1 was not changed upon phosphatase treatment (Fig. S2A) and it has been reported to be N-glycosylated(*42*), we performed immunoblotting of total lysates prepared from Ypf1-FLAG-expressing cells with or without Endo H treatment. The results showed that Ypf1 appeared as a doublet under both glucose-rich and -depleted conditions, with the upper band exhibiting a stronger signal during GS. Upon Endo H treatment, both bands merged into a single band with a lower molecular weight than the original bands whereas Nvj1 and Nsg2 were unaffected (Fig. 2F, S2B). The molecular weight of Endo H- treated Ypf1 was identical to that of the Ypf1-3NQ mutant, in which all potential N-glycosylation sites (N22, N23, and N28) in the N-terminal region were substituted with glutamine. These findings indicate that the N-terminal region of Ypf1 undergoes additional N-glycosylation in a GS-dependent manner. To identify the glycosylation site, we generated Ypf1-N22Q, N23Q, and N28Q mutants and analyzed their migration patterns. We found that GS-dependent glycosylation was abolished in the N28Q mutant, demonstrating that Asn28 is specifically glycosylated in response to GS. This suggests that under normal conditions, Asn28 is not exposed to the luminal space where glycosylation can occur, whereas under glucose-starved condition, it becomes accessible and undergoes glycosylation, implying a GS-induced structural rearrangement in the N-terminal region of Ypf1. The N22Q mutation partially suppressed glycosylation under nutrient-rich conditions, but not under glucose-starved conditions (Fig. 2G). The N23Q mutation did not affect the migration pattern of Ypf1 under both normal and glucose-starved conditions (Fig. 2G). Given that the 3NQ mutations completely abolish glycosylation (Fig. 2F), Asn22 is the primary site for glycosylation while Asn23 can also be glycosylated. Furthermore, we confirmed that the absence of Ypf1, Nsg1, or Nsg2 did not affect the steady-state levels or modification patterns of Nvj1, Ypf1, Nsg1, and Nsg2 under GS condition (Fig. 2H).

### Ypf1 and Snf1 mediate NVJ remodeling during GS

Among the GS-dependent NVJ factors, we found that Ypf1 played a central role in recruiting other components to the NVJ. Loss of Ypf1 impaired the NVJ partitioning of Nsg1, Nsg2, Hmg1, and Hmg2 but not that of Nvj1, Nvj2, Vac8, Osh1, and Tsc13 (Fig. 3A). In contrast, Nsg1 and Nsg2 were not required for the NVJ partitioning of Ypf1 and Hmg1 (Fig. 3B). The glycosylation and peptidase activity of Ypf1 were dispensable for the NVJ partitioning of both Ypf1 and Hmg1, as the glycosylation-deficient mutant Ypf1-3NQ and the catalytically inactive mutant Ypf1-D366,411A showed localization patterns comparable to the wild-type (Fig. S3A-C) The loss of Ypf1 did not affect the steady-state levels or post-translational modification patterns of Nvj1, Nsg1, and Nsg2 (Fig. 3C). Together, these results strongly suggest that Ypf1 functions as a GS sensor that recruits Nsg1, Nsg2, Hmg1, and Hmg2 to the NVJ.

**Fig. 3.**
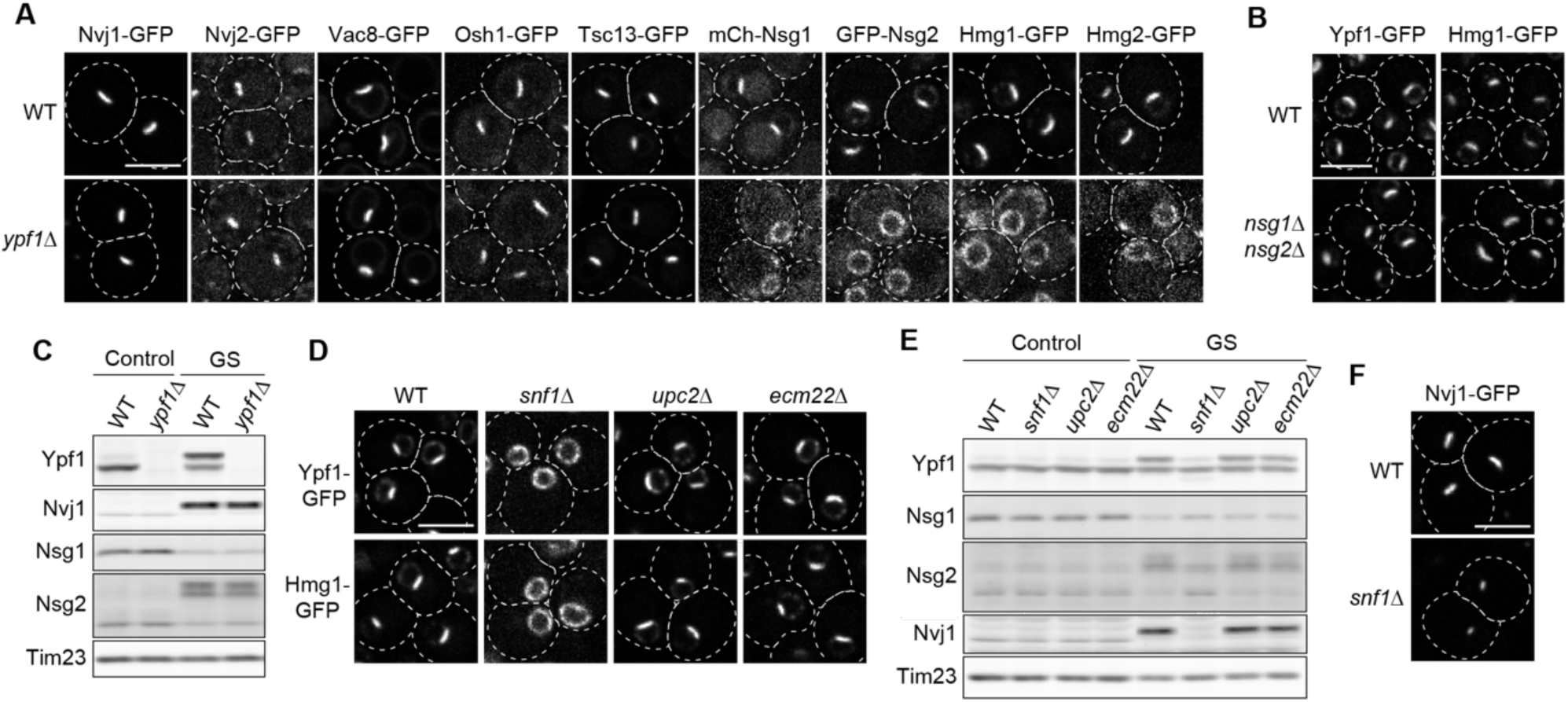
GS-dependent NVJ partitioning depends on Ypf1 and Snf1. (**A**) Wild-type and ypf1Δ cells expressing the indicated NVJ proteins fused to GFP or mCherry were observed by confocal fluorescence microscopy after a 24-hour GS treatment. (**B**) Wild-type and nsg1Δnsg2Δ cells expressing Ypf1-GFP or Hmg1-GFP were imaged by confocal fluorescence microscopy after a 24-hour GS treatment. (**C**) Immunoblotting of whole cell extracts prepared from WT and ypf1Δ cells grown with or without a 24-hour GS treatment. (**D**) The indicated cells expressing the Ypf1-GFP or Hmg1-GFP were imaged by confocal fluorescence microscopy after a 24-hour GS treatment. Images in (**A**), (**B**), and (**D**) are single focal plane. (**E**) Immunoblotting of whole cell extracts prepared from the indicated cells grown with or without a 24-hour GS treatment. (**F**) Wild-type and snf1Δ cells expressing Nvj1-GFP were observed by confocal fluorescence microscopy after a 24-hour GS treatment. Maximum projection images reconstituted from z-stacks are shown. Scale bars, 5 µm.

We next investigated upstream regulators of Ypf1. Previous studies reported that the ergosterol-responsive transcription factor Upc2 (*43*) is important for the localization of Hmg1 to the NVJ (*31*). However, our experiments showed that the deletion of *UPC2* or its paralog *ECM22* (*43*) did not affect the NVJ localization of Ypf1 and Hmg1 (Fig. 3D). In contrast, deletion of Snf1, an AMP-activated S/T protein kinase (AMPK) that is important for glucose-responsive transcriptional regulation (*44*), prevented the NVJ partitioning of Ypf1 and Hmg1 (Fig. 3D). Moreover, Snf1 deficiency markedly impaired the GS-dependent glycosylation of Ypf1, reduced the phosphorylation levels of Nvj1 and Nsg2, and destabilized Nvj1. Based on these results, two possible mechanisms could explain why Ypf1 and Hmg1 fail to localize to the NVJ in the absence of Snf1. One possibility is that the GS-dependent structural change of Ypf1, as indicated by its glycosylation, does not occur without Snf1. Another possibility is that reduced Nvj1 levels caused by Snf1 deletion impair the NVJ partitioning of Ypf1 and Hmg1. However, we confirmed that Nvj1 remains localized to the NVJ in *snf1*Δ cells (Fig. 3F). We also observed that the GS-induced increase in Nvj1 levels was significantly suppressed when GFP was fused to Nvj1 (Fig. S3D). Nevertheless, Ypf1 and Hmg1 were still able to localize to the NVJ in cells expressing Nvj1- GFP (Fig. 2A). These findings suggest that the NVJ localization of Nsg1, Nsg2, Hmg1, and Hmg2 is primarily impaired by the absence of GS-induced structural changes in Ypf1, rather than by reduced Nvj1 levels due to Snf1 deficiency.

### Ypf1 senses GS through VLCFA metabolism changes

What does Ypf1 sense during GS to trigger NVJ remodeling? Since Asn28 of Ypf1 is glycosylated in a GS-dependent manner, it is likely that the N-terminal region of Ypf1 undergoes a conformational change. We hypothesized that alterations in the lipid composition of the nuclear membrane during GS may alter the membrane topology of the N-terminal region of Ypf1. To test this hypothesis, we performed a lipid-focused screen to identify genes whose deletion affects Ypf1 glycosylation. Specifically, we analyzed the GS-dependent glycosylation of Ypf1 in yeast strains lacking genes involved in phospholipid, sphingolipid, ergosterol, or fatty acid metabolism. Ypf1 glycosylation was enhanced under both nutrient-rich and glucose-starved conditions in cells lacking *ELO3*, which encodes a fatty acid elongase responsible for the synthesis of very long-chain fatty acids (VLCFAs) (*45*) (Fig. S4A). We therefore examined whether loss of Elo3 also influenced other GS-dependent NVJ phenotypes. Intriguingly, Elo3 deficiency markedly promoted the phosphorylation and stabilization of Nvj1 and Nsg2, as well as the destabilization of Nsg1 (Fig. 4A). In addition, NVJ partitioning of Ypf1 was accelerated and enhanced in *elo3*Δ cells (Fig. 4B). These findings suggest that depletion of VLCFA synthesis facilitates GS-induced NVJ remodeling.

**Fig. 4.**
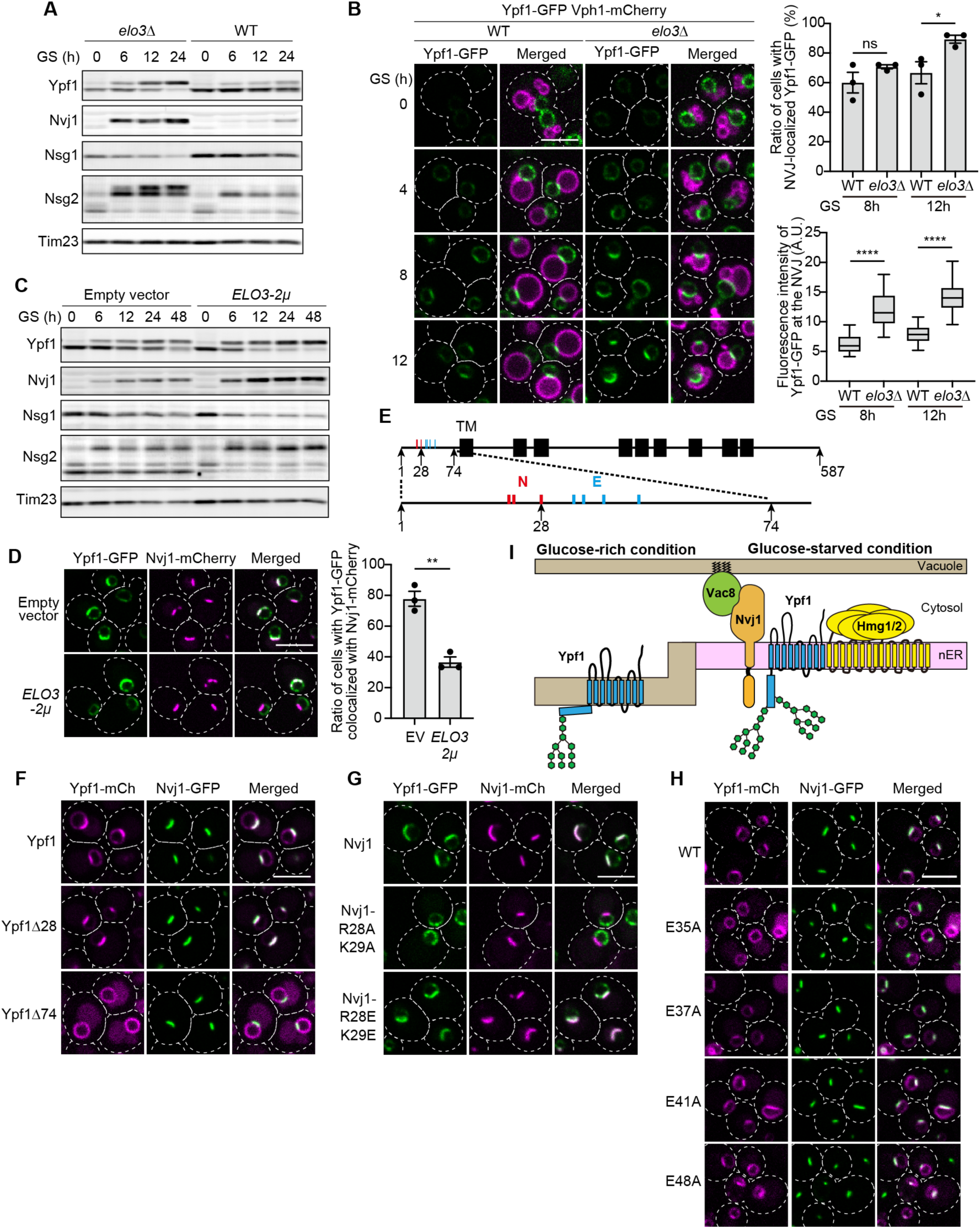
VLCFA biogenesis is involved in NVJ remodeling. (**A**) Immunoblotting of wild-type and elo3Δ cell extracts after GS treatments for the indicated durations. (**B**) The progression of Ypf1-GFP NVJ partitioning was assessed in cells expressing Vph1-mCherry after GS treatment for the indicated time periods. Single focal plane images obtained by confocal fluorescence microscopy are shown. Bar graph shows the percentage of cells displaying NVJ-localized Ypf1-GFP. Values are means ± S.E. (n = 3). At least 25 cells were analyzed per biological replicate. Box-and-whisker plot shows the fluorescence intensity of Ypf1-GFP at the NVJ (n = 25 cells). ns: not significant, ∗: p< 0.05, ∗∗∗∗: p< 0.0001. p-values were obtained from the unpaired two-tailed t-test. (**C**) Immunoblotting of whole-cell extracts prepared from cells carrying either an empty vector or a multi-copy plasmid encoding the ELO3 gene, following GS treatment for the indicated time periods. (**D**) Yeast cells expressing Ypf1-GFP and Nvj1-mCherry and carrying either an empty vector or a 2 μ plasmid encoding ELO3 were observed by confocal fluorescence microscopy after 1 day of GS treatment. Bar graph shows the percentage of cells with Ypf1-GFP localized to the NVJ marked by Nvj1-mCherry. Values are means ± S.E. (n = 3). At least 30 cells were analyzed per biological replicate. ∗∗: p< 0.01. p-values were obtained using the unpaired two-tailed t-test. **(E**) A schematic diagram of Ypf1 showing its transmembrane domains (black boxes). Red lines indicate asparagine residues that undergo N-glycosylation, and blue lines indicate glutamic acid residues. (**F**, **G**, **H**) Yeast cells expressing C-terminally GFP or mCherry-fused Ypf1 and Nvj1 mutants were imaged by confocal fluorescence microscopy after 1 day of GS. All images shown in (**B**), (**D**), (**F**), (**G**), and (**H**) are single focal plane. Scale bar, 5 μm. (**I**) Schematic model of the NVJ remodeling under GS. Color differences in the nuclear ER (nER) indicate predicted changes in membrane properties.

We then asked whether overexpression of Elo3 would have the opposite effect. Expression of Elo3 from the *2µ* multi-copy plasmid slightly enhanced the GS-dependent glycosylation of Ypf1, as well as phosphorylation and stabilization of Nvj1 and Nsg2, and destabilization of Nsg1 (Fig. 4C). Nevertheless, NVJ partitioning of Ypf1 was inhibited in cells overexpressing Elo3 (Fig. 4D). These results suggest that imbalances in VLCFA metabolism affect GS-dependent NVJ factor dynamics, however, suppression of VLCFA synthesis or Elo3 activity is crucial for proper progression of NVJ remodeling during GS.

We next asked whether the N-terminal region of Ypf1, which likely undergoes changes in membrane orientation, was critical for the GS-dependent NVJ partitioning. To this end, we created truncated mutants, Ypf1Δ28 and Ypf1Δ74, lacking its N-terminal 28 and 74, amino acids, respectively, and observed their localization (Fig. 4E). The results showed that Ypf1Δ74, but not Ypf1Δ28, failed to localize to the NVJ during GS, suggesting that amino acids 29-74 of Ypf1 are essential for its NVJ partitioning to the NVJ (Fig. 4F). We next examined how the N-terminal region contributes to NVJ remodeling. Previous studies reported that the positively charged residues R28 and K29 in the luminal domain of Nvj1 are essential for Hmg1 localization to the NVJ (*31*), raising the possibility that other GS-specific NVJ factors might also depend on these residues. We therefore examined the localization of Ypf1, Nsg1, and Nsg2 in the Nvj1-R28A/K29A mutant cells (Fig. 4G and S4B). None of these proteins localized to the NVJ in the mutants, suggesting that Ypf1 is recruited to the NVJ through a direct electrostatic interaction between acidic residues in its N-terminal region and positively charged residues of Nvj1. However, this hypothesis was not supported by further analysis. Ypf1 still accumulated at the NVJ in Nvj1-R28E/K29E mutant cells, in which the charge is inverted (Fig. 4G). We also examined Ypf1 mutants with alanine substitutions at individual acidic residues in the N-terminal region. Ypf1-E37A, -E41A, and -E48A showed localization similar to the wild-type, whereas only Ypf1-E35A failed to accumulate at the NVJ during GS (Fig. 4H). These results suggest that both the luminal domains of Nvj1 and Ypf1 are important for NVJ remodeling, but that they may not interact through direct electrostatic interactions. This observation raises the possibility that an as-yet-unidentified factor mediates the interaction between Nvj1 and Ypf1 during GS.

### Nsg1/Nsg2 loss causes lipid droplet accumulation during GS

We next sought to address the physiological role of GS-dependent NVJ remodeling. Previously, the GS-dependent Hmg1 NVJ partitioning was reported to activate mevalonate pathway flux. In addition, Nsg1 and Nsg2 have been reported to be important for stability of Hmg2 (*32*, *46*, *47*). However, the significance of reciprocal effects on the stabilities of Nsg1 and Nsg2 remains unknown. We thus created yeast strains lacking Nsg1, Nsg2, Hmg1, and Hmg2 and examined their phenotypes. We first noted the accumulation of lipid droplet (LD)-like spherical structures in DIC images of *nsg1Δnsg2Δ* cells (Fig. 5A). LipiBlue, a dye that stains lipid droplets (LDs), and LD-marker Erg6-GFP labeled these spherical structures, confirming that they are LDs (Fig. 5A, S5). In wild-type, *nsg1*Δ, *nsg2Δ*, and *nsg1Δnsg2Δ* cells, approximately 20% of the cells contained LDs that were strongly stained with LipiBlue, while the remaining 80% harbored only weakly stained LDs (Fig. 5B). In *hmg1Δnsg1Δnsg2Δ* and *hmg2Δnsg1Δnsg2Δ* cells, the proportion of cells with strongly stained LDs was slightly increased (Fig. 5B). These differences in staining intensity with LipiBlue suggest variations in the lipid composition of the LDs. Regardless of whether the cells contained strongly or only weakly stained LDs, a pronounced accumulation of LDs was observed in *nsg1Δnsg2Δ* cells (Fig. 5C). To examine whether LD accumulation was due to defective lipophagy, we observed LDs in cells lacking Atg1, a core component of macroautophagy that also plays a role in lipophagy(*48–51*). LDs did not accumulate in *atg1*Δ cells, as in wild-type cells, indicating that the LD accumulation seen in *nsg1Δnsg2Δ* cells is not attributable to impaired lipophagy. (Fig. 5A, C). Interestingly, the LD accumulation observed in *nsg1Δnsg2Δ* cells was abolished by deletion of *HMG1*, but not by deletion of *HMG2*, highlighting the importance of Hmg1 activity for LD accumulation in *nsg1Δnsg2Δ* cells.

**Fig. 5.**
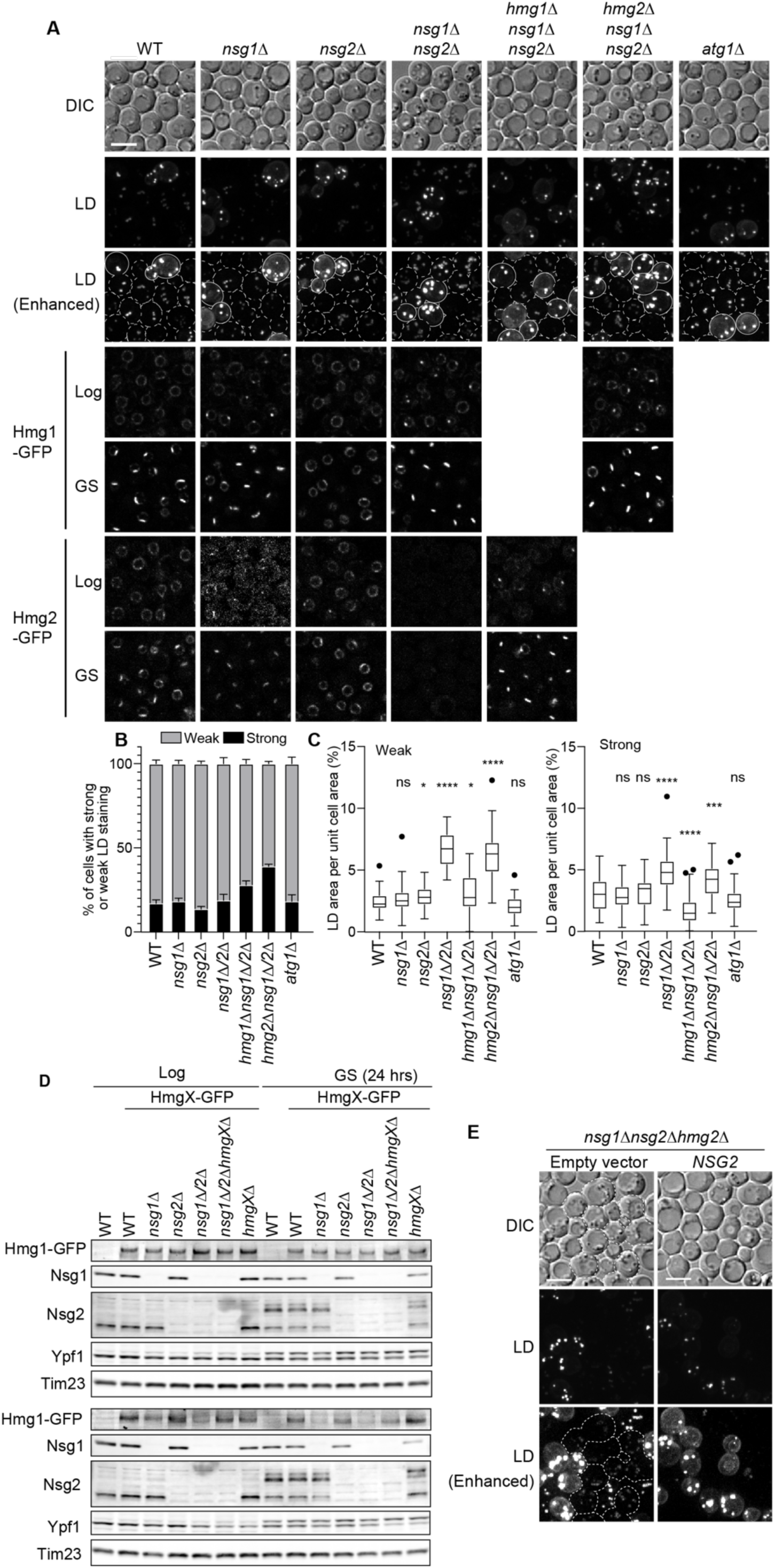
LDs accumulate in an Hmg1-dependent manner in *nsg1Δnsg2Δ* cells. (**A**) LDs in the indicated cells after a 1-day GS treatment were visualized using the LD- staining dye LipiBlue and imaged by confocal fluorescence microscopy. The indicated yeast cells expressing Hmg1-GFP or Hmg2-GFP were observed by confocal fluorescence microscopy under glucose-rich (Log) and glucose-starved (GS) conditions. Yeast cells with strongly and weakly stained LDs were outlined with solid and dotted lines, respectively. LDs are shown as maximum intensity projection images. Fluorescence images of Hmg1-GFP and Hmg2-GFP represent a single optical section. Scale bars, 5 µm. (**B**) Bar graph shows the populations of cells with weak and strong LD fluorescence intensity based on maximum projection images acquired in (**A**). Values are means ± S.E. (n = 4). At least 100 cells were analyzed per biological replicate. (**C**) Box-and-whisker plot shows the percentage of LD area relative to total cell area (n = 40 cells). ns: not significant, ∗: p< 0.05, ∗∗: p< 0.01, ∗∗∗: p< 0.001 ∗∗∗∗: p< 0.0001. p-values were obtained using the unpaired two-tailed t-test. (**D**) Whole-cell lysates prepared from the indicated yeast cells, with or without a 1-day GS treatment, were analyzed by immunoblotting. HmgX denotes either Hmg1 or Hmg2. (**E**) LDs in nsg1Δnsg2Δ cells carrying either an empty vector or a CEN-plasmid expressing Nsg2, were stained with LipiBlue and visualized by confocal fluorescence microscopy after a 1-day GS treatment. Yeast cells showing weak LD staining were outlined with dotted lines. LDs are shown as maximum intensity projection images. Scale bars, 5 µm.

We then examined Hmg1 localization in these gene-deletion mutant cells. Hmg1 did not accumulate at the NVJ in wild-type cells under glucose-rich conditions. However, in the absence of Nsg1, Hmg1 partly relocated to the NVJ even under glucose-rich conditions (Fig. 5A, *nsg1*Δ, *nsg1Δnsg2*Δ, and *nsg1Δnsg2Δhmg2Δ*). Under the GS condition, the Hmg1 NVJ partitioning was partially compromised in *nsg2*Δ cells, whereas it was restored when Nsg1 was additionally deleted (Fig. 5A, *nsg1Δsng2Δ*). As shown in Fig. 2E, Nsg1 is stably present under glucose-rich conditions but decreases dramatically during GS. These findings suggest that Nsg1 suppresses Hmg1 NVJ partitioning in the presence of glucose, while the GS-dependent destabilization of Nsg1 allows Hmg1 to accumulate at the NVJ for Hmg1 activation. However, we observed LD accumulation in *nsg1*Δ*nsg*2Δ but not *nsg1Δ* cells, indicating that enhanced Hmg1 NVJ partitioning is not the sole factor responsible for LD accumulation.

We also observed the localization of Hmg2, another HMG-CoA reductase. Similar to Hmg1, Hmg2 was not localized to the NVJ under glucose-rich conditions, whereas it accumulated there under GS conditions. In *nsg1Δ* cells, Hmg2-GFP relocated to the NVJ under GS as in wild-type cells, although the amount of Hmg2-GFP was lower, as the stability of Hmg2 is dependent on Nsg1(*32*) (Fig. 5A, D). Similar to the case of Hmg1-GFP, the NVJ partitioning of Hmg2-GFP was partially impaired in *nsg2*Δ cells. As the loss of Nsg1and Ngs2 destabilized Hmg2 (Fig. 5D), we no longer detected Hmg2-GFP signal in *nsg1Δnsg2Δ* cells. In contrast, loss of Hmg1 partly restored the decreased levels of Hmg2- GFP, enabling us to observe clear NVJ partitioning of Hmg2-GFP in *nsg1*Δ*nsg*2Δ*hmg1Δ* cells (Fig. 5A and 5D). This suggests that Hmg1 negatively regulates the stability and NVJ partitioning of Hmg2. Taken together, these results indicate that LD accumulation occurs only when Nsg1 and either Nsg2 or Hmg2 are non-functional.

We thus asked whether Nsg2 or Hmg2 deficiency, when combined with the loss of Nsg1, leads to LD accumulation. To address this, we introduced a *CEN* plasmid expressing Nsg2 into *nsg1Δnsg2Δhmg2Δ* and observed LDs. The result showed that re-expression of Nsg2 in *nsg1Δnsg2Δhmg2Δ* cells markedly reduced the number of cells containing LDs that are only weakly stained by LipiBlue (Fig. 5E). This result was also confirmed by DIC images, which showed a clear reduction in LD accumulation upon Nsg2 expression (Fig. 5E, dotted line). These results indicate that the loss of Nsg2, but not Hmg2, in addition to the absence of Nsg1, cause the LD accumulation. These findings strongly suggest that both Nsg1 and Nsg2 not only play a role in stabilizing Hmg2, but also function as suppressors of Hmg1 activity. Specifically, in the presence of glucose, Nsg1 remains stable, preventing Hmg1 NVJ partitioning, while the expression level of Nsg2 is kept low, maintaining basal levels of HMG-CoA reductase activity. During GS, Nsg1 becomes unstable, promoting Hmg1 accumulation at the NVJ and activating its activity. Conversely, GS stabilizes Nsg2, which could in turn suppress HMG-CoA reductase activity. Together, these findings support the idea that Nsg2 contributes to a negative feedback mechanism that limits ergosterol synthesis and maintains sterol homeostasis during GS.

### Loss of Nsg1 and Nsg2 drives excessive ergosterol synthesis

To test the idea above that Nsg1 and Nsg2 regulate the HMG-CoA reductase activity as suppressors, we investigated ergosterol synthesis. Specifically, we metabolically labeled yeast cells with ^14^C-acetate and analyzed radioisotope (RI)-labeled sterol lipids by thin-layer chromatography. As a control, we also analyzed phospholipids using non-saponified samples, and normalized levels of sterol and its precursor lipids to phospholipid content for comparison. Under glucose-rich conditions, there was no significant difference in RI- labeled sterol and its precursor lipid levels between wild-type and *nsg1Δnsg2Δ* cells (Fig. 6A, B). However, intriguingly, under GS, ergosterol esters and squalene, a triterpene intermediate in sterol biosynthesis were drastically accumulated in *nsg1Δnsg2Δ* cells, strongly suggesting hyperactivation of HMG-CoA reductase in the absence of Nsg1 and Nsg2 (Fig. 6A, B). The drastic increase in ergosterol ester and squalene was also observed in *nsg1Δnsg2Δhmg2Δ* cells, which exhibit abnormal lipid droplet accumulation. In contrast, we did not observe such lipid accumulation in *nsg1Δnsg2Δhmg1Δ* cells, where lipid droplet accumulation was not detected. Although the loss of Nsg1 or Nsg2 alone did not drastically affect sterol synthesis, we observed a slight inhibition in sterol synthesis in *nsg1*Δ cells and a slight increase in *nsg2*Δ mutants. These findings suggest that when both Nsg1 and Nsg2 are lacking, Hmg1 becomes hyperactive, leading to the excessive accumulation of ergosterol esters and squalene, which in turn drives abnormal LD formation. Thus, we propose that the INSIG proteins Nsg1 and Nsg2 are not only involved in regulating Hmg2 stability but also play a critical role in repressing Hmg1 activity (Fig. 7). This Hmg1 suppression may constitute a negative feedback mechanism that prevents excessive sterol accumulation under GS conditions.

**Fig. 6.**
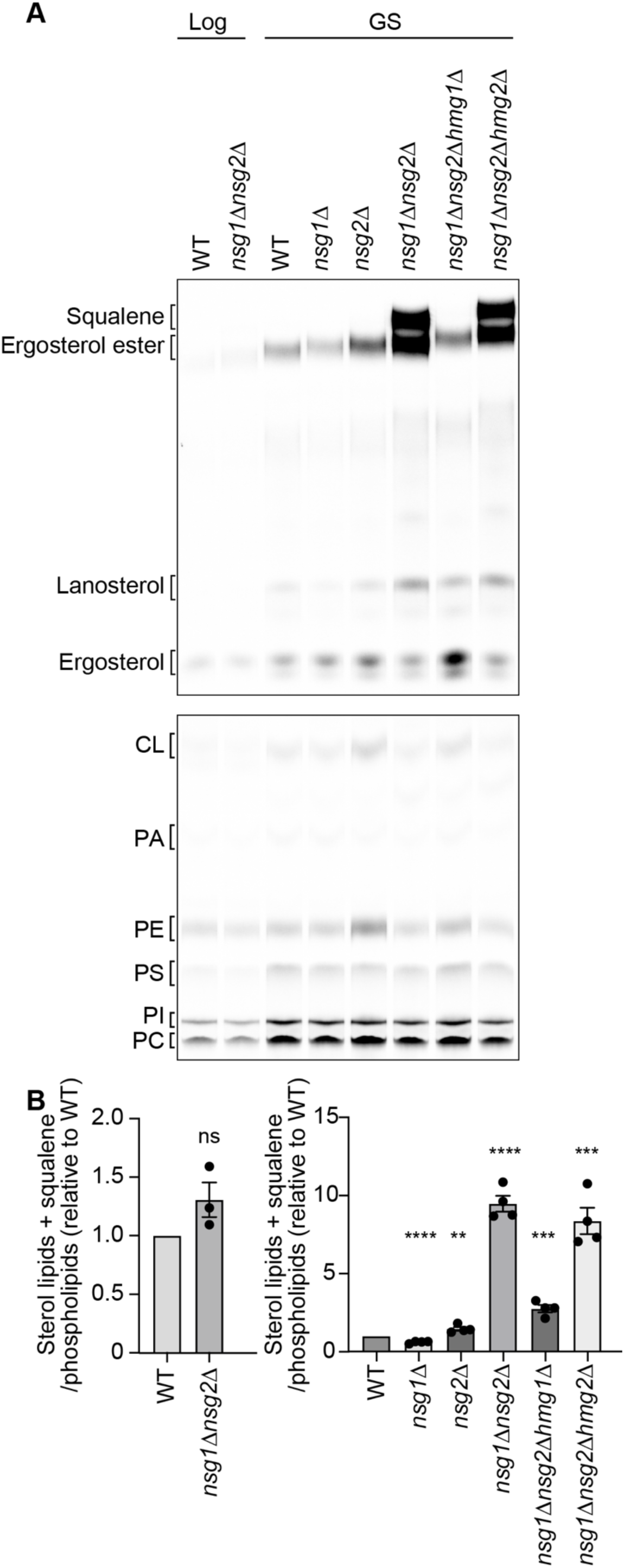
Squalene and ergosterol esters accumulate in an Hmg1-dependent manner in *nsg1Δnsg2Δ* cells. (**A**) The indicated yeast cells were cultured for 1 day in YPD or GS medium containing ¹⁴C-acetate, after which total lipids were extracted. The extracted lipids were treated with or without saponification, separated by thin-layer chromatography, and visualized by autoradiography. (**B**) Bar graph shows the combined levels of sterol lipids and squalene, normalized to phospholipid content. Wild-type value was set to 1. Values are means ± S.E. (n = 3). ns: not significant, ∗∗: p< 0.01, ∗∗∗: p< 0.001 ∗∗∗∗: p< 0.0001. p -values were obtained using the unpaired two-tailed t-test.

**Fig. 7.**
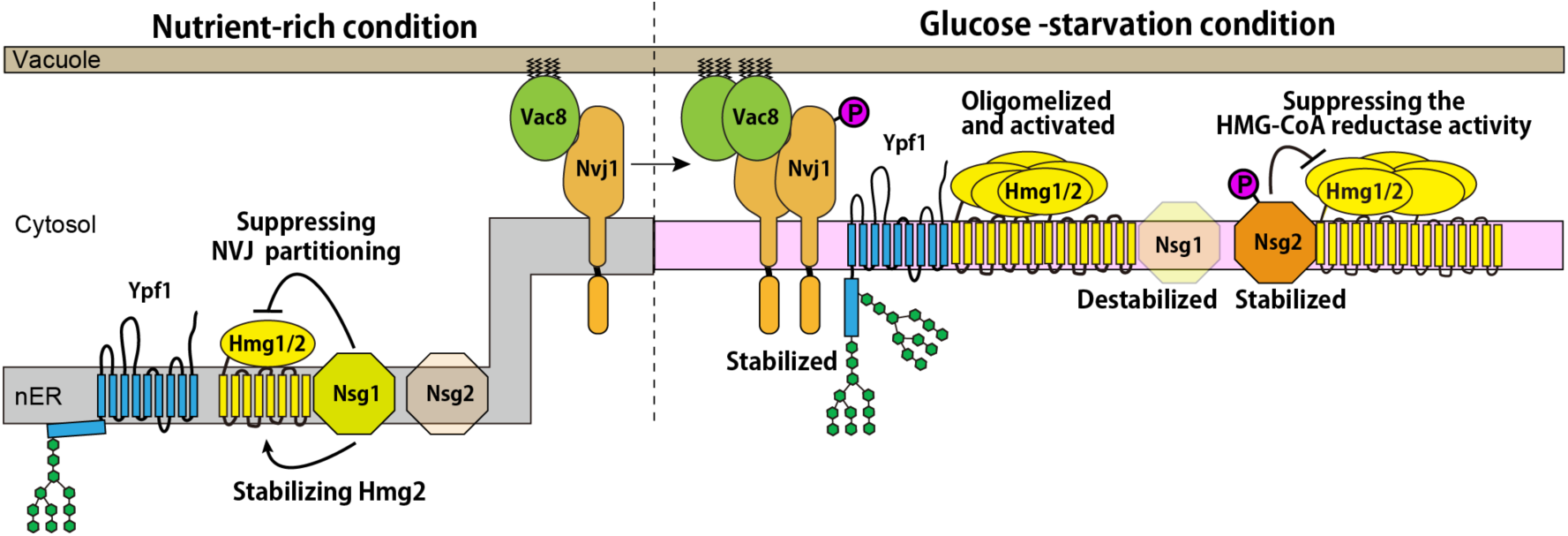
A model for GS-dependent NVJ remodeling. Schematic model of NVJ remodeling under GS conditions. Color differences in the nuclear ER (nER) indicate predicted changes in membrane properties.

In summary, our findings reveal a novel stress-response mechanism in which GS triggers membrane remodeling of the NVJ through changes in VLCFA-containing lipid levels, ensuring the proper regulation of Hmg1 activity (Fig. 7). This insight significantly advances our understanding of the role of the NVJ, as well as VLCFAs in cellular stress responses. Moreover, beyond its fundamental biological implications, this study provides a foundation for the development of yeast-based strategies to enhance the industrial production of commercially valuable lipids such as ergosterol and squalene.

## Discussion

MCSs are dynamic inter-organelle interfaces that play central roles in coordinating lipid metabolism and stress adaptation. Previous studies have highlighted the structural plasticity of MCSs such as ERMES, vCLAMP, and NVJ (*9–12*, *16*, *30*, *31*, *35*). However, the regulatory mechanisms and physiological roles of MCS remodeling have remained poorly understood. In this study, we focused on the NVJ and demonstrate that NVJ remodeling is tightly linked to VLCFA metabolism and is critical for regulating ergosterol biosynthesis in response to GS (Fig. 7).

We identified the INSIG homologs Nsg1 and Nsg2, along with the aspartyl protease Ypf1, as NVJ-localized factors that accumulate in a GS-dependent manner (Fig. 2). Our results show that Ypf1 promotes the recruitment of Nsg1, Nsg2, Hmg1, and Hmg2 to the NVJ under GS (Fig. 3), and that Nsg1 and Nsg2 act as negative regulators of Hmg1 (Fig. 5 and 6). These two INSIG homologs exhibit opposing stabilities during GS: Nsg1 is destabilized, while Nsg2 is stabilized (Fig. 2E), allowing for dynamic tuning of Hmg1 activity likely through a negative feedback mechanism that maintains appropriate ergosterol levels. Consistent with this idea, simultaneous deletion of Nsg1 and Nsg2 led to hyperactivation of Hmg1 and excessive accumulation of ergosterol esters and squalene (Fig. 6).

Our data suggest that these regulatory events are driven by GS-dependent suppression of VLCFA synthesis. Although Elo3 deletion clearly promotes NVJ phenotypes that occur during GS (Fig. 4), the precise biophysical consequences of VLCFA depletion or Elo3 deficiency remain to be clarified. It is currently unknown whether the loss of Elo3 leads to decreased membrane thickness or alters other membrane properties such as fluidity, tension, or curvature. A recent study reported that proteins with short transmembrane domains preferentially accumulate at the NVJ under GS conditions (*52*). This observation highlights the potential role of membrane thickness in NVJ-specific protein recruitment and also offers a new perspective on the well-known localization of Tsc13, an enoyl reductase that catalyzes the last step in each cycle of VLCFA elongation, at the NVJ. While Tsc13 is known to be required for PMN (*25*), the physiological significance of its localization at the NVJ beyond this role has remained unclear. Our results suggest that local VLCFA synthesis at the NVJ contributes to defining membrane properties essential for the spatial recruitment of NVJ regulatory proteins in response to GS. Supporting this idea, Tsc13 is reported to be coimmunoprecipitated with Elo2 and Elo3 (*21*, *53*). This coordination between lipid synthesis and protein recruitment provides a mechanistic framework for understanding how NVJ remodeling adapts to cellular metabolic demands.

Despite these advances, important mechanistic questions remain. Most notably, how Ypf1 recruits Nsg1, Nsg2, and Hmg1/2 to the NVJ remains unclear. We found that the luminal domains of both Nvj1 and Ypf1 are critical for NVJ localization (Fig. 4), but direct physical interaction between these proteins was not detected (data not shown). This suggests the existence of yet-to-be-identified bridging factors that may interact with the N-terminal domains of Nvj1 and Ypf1. Identification of such factors will be key to elucidating the molecular basis of NVJ remodeling. Besides, the upstream mechanism by which VLCFA synthesis is suppressed during GS remains unclear. It is not yet known whether this suppression is mediated by a reduction in Elo3 levels, potentially through transcriptional, translational, or post-translational regulation. Elucidating how VLCFA metabolism is downregulated in response to GS will be critical for fully understanding the link between membrane composition and metabolic stress adaptation.

We also found that Ypf1 likely undergoes structural changes during GS, as indicated by alterations in its N-glycosylation pattern (Fig. 2). While Ypf1 is conserved from yeast to humans, its N-terminal region is poorly conserved, implying that this GS-responsive remodeling mechanism may be specific to yeast. Nevertheless, regulation of membrane physical properties via VLCFA metabolism is likely relevant across species. In mammalian cells, for instance, late endosome/lysosomes (which functionally parallel yeast vacuoles) are known to extensively interact with the ER, forming ER-lysosome MCSs (*54*). Investigating whether VLCFA-dependent changes in membrane composition regulate factor recruitment at these sites may uncover conserved MCS-based stress response mechanisms. Moreover, if the N-terminal domain of Ypf1 truly undergoes conformational changes in response to membrane properties, it may serve as a foundation for developing novel membrane-responsive biosensors.

Additional mechanistic questions also remain regarding the regulation of Nsg1 and Nsg2 stability. Previous studies have shown that Nsg1 stability depends on lanosterol availability and that it is degraded in the absence of this sterol intermediate (*46*). Given that GS stimulates ergosterol biosynthesis, the resulting depletion of lanosterol may contribute to Nsg1 destabilization. Conversely, Nsg2 stabilization during GS may be mediated by phosphorylation. Future analysis using phospho-deficient and phospho-mimetic mutants will be important to elucidate this regulatory mechanism. Furthermore, we observed that Hmg2 becomes stabilized upon Hmg1 deletion, even in cells lacking both Nsg1 and Nsg2 (Fig. 5A, D). This suggests that additional, unidentified factors may regulate Hmg2 turnover, independent of the known INSIG homologs.

Beyond its fundamental biological significance, our findings have important implications for biotechnology. For example, squalene, a valuable lipid used as an adjuvant, in cosmetics, and in dietary supplements, is still largely sourced from deep-sea shark liver oil (*55*). Due to ethical and sustainability concerns, there is growing demand for alternative, non-animal production systems (*55*, *56*). Although metabolic engineering strategies using yeast have achieved some success in boosting squalene production, primarily by enhancing acetyl-CoA supply (*57*) or overexpressing sterol biosynthetic enzymes (*58*, *59*), these efforts often overlook endogenous negative feedback mechanisms that limit sterol accumulation. Our study reveals one such feedback circuit involving Nsg1 and Nsg2. Disrupting this regulatory loop could synergize with existing metabolic flux optimization strategies to achieve greater production yields. If the repression of this feedback circuit enables more efficient synthesis of squalene or its precursors such as farnesene, it could pave the way for more sustainable and ethically acceptable squalene production platforms.

## Materials and Methods

### Yeast strains and growth conditions

*Saccharomyces cerevisiae* strain FY833 (MAT*a ura3-52 his3-Δ200 leu2-Δ1 lys2-Δ202 trp1-Δ63*) was used as background strains (*60*). The yeast cells used in this study are listed in Table S1. Yeast knockout strains used to screen for genes whose deletion affects the glycosylation pattern of Ypf1 are listed in Table S2. The C-terminal tagging, and gene disruptions were performed by homologous recombination using the appropriate gene cassettes amplified from the plasmids listed in Data S1 (*61*). The primer pairs used in this study was summarized in Table S3. To introduce the GFP or mCherry tag for the N-terminus of *NSG1* and *NSG2*, CRISPR-Cas9 system was used as described previously (*62*). Briefly, we first selected guide RNA target sequences around the start codons of *NSG1* and *NSG2* using CRISPRdirect. After hybridizing the pair of oligonucleotides containing the target sequences (YU5281/YU5282 for *NSG1* and YU5286/YU5287 for *NSG2*), they were introduced into the Cas9 expression plasmid 16-15 (pYU207), which had been digested with BsaI. The resulting plasmids were introduced into yeast cells along with donor DNA fragments encoding GFP or mCherry, which were flanked by sequences homologous to the regions 50 bp upstream and downstream of the start codons of *NSG1* and *NSG2*. The resulting transformants were cultured in SCGal-Ura medium, and the integration of the desired DNA fragments was confirmed by PCR using genomic DNA as a template. Finally, the Cas9 expression plasmid was eliminated by culturing the cells in SCD+FOA medium.

Yeast strains chromosomally expressing CsFiND proteins or TurboID-HA were generated as previously described. Briefly, DNA fragments encoding Ifa38-GFP(n)- 3xFLAG-TurboID(c)-natNT2, Dpp1-TurboID(n)-V5-GFP(c)-hphMX, or TurboID-HA- hphMX were PCR-amplified with flanking sequences homologous to the *URA3* or *LEU2* loci and integrated into the corresponding genomic regions. All plasmids used in this study and their construction procedures are summarized in Data S1.

Yeast cells were cultured in YPD (1% (w/v) yeast extract, 2% (w/v) polypeptone, and 2% glucose), SCD (0.67% (w/v) yeast nitrogen base without amino acids, 0.5% (w/v) casamino acids, and 2% (w/v) glucose) or SCGal (0.67% (w/v) yeast nitrogen base without amino acids, 0.5% (w/v) casamino acids, and 2% (w/v) galactose) with appropriate supplements. To eliminate *URA3*-containing plasmids, 5-Fluoroorotic acid was added to SCD medium at a final concentration of 1 µg/ml. To induce glucose starvation, YP or SCD medium containing 0.01% (w/v) glucose was used. For nitrogen starvation, cells were cultured in SD-N medium containing 0.17% (w/v) yeast nitrogen base without amino acids and ammonium sulfate, and 2% (w/v) glucose.

### Antibodies

Polyclonal antibodies against Nvj1, Nvj2, Vac8, Ypf1, Nsg1, and Nsg2 were generated by immunizing rabbits with N-terminally His-tagged recombinant proteins expressed in *Escherichia coli*. The antigens corresponded to the following amino acid regions: Nvj1 (residues 229–321), Nvj2 (residues 218–492), Vac8 (residues 10–515), Ypf1 (residues 497– 587), Nsg1 (residues 1–90), and Nsg2 (residues 1–100). Among these, only Nsg1 was purified as a soluble protein, while the other recombinant proteins were isolated from inclusion bodies.

### Purification of Biotinylated Proteins by CsFiND

Yeast cells expressing CsFiND proteins were inoculated into 200 ml of SCD at an initial OD₆₀₀ of 0.02. Once the OD₆₀₀ reached between 0.8 and 1.2, biotin was added to a final concentration of 50 μM, followed by an additional 3-hour incubation. To induce nitrogen starvation, logarithmically growing cells were collected and washed with SD-N medium twice and transferred to SD-N medium for 3 h. Biotin was then added to a final concentration of 50 μM, followed by an additional 3-hour incubation. Cells corresponding to 200 OD₆₀₀ units were harvested and incubated in 20 ml of alkaline buffer (0.1 M Tris-HCl, pH 9.5, 10 mM DTT) at 30°C for 10 min. After washing with spheroplast buffer (20 mM Tris-HCl, pH 7.5, 1.2 M sorbitol), the cells were treated with 2 units/ml of Zymolyase 20T (Nacalai Tesque, Inc.) in 25 ml of spheroplast buffer and incubated for 30 min at 30°C. The resulting spheroplasts were washed with ice-cold spheroplast buffer and lysed by vortexing for 1 minute in 1 ml of ice-cold breaking buffer (20 mM Tris-HCl, pH 7.5, 0.6 M mannitol, 1 mM EDTA, 1 mM PMSF) containing 0.5 ml of glass beads. An additional 5 ml of breaking buffer was added, and the mixture was centrifuged at 2,000 × *g* for 5 min to remove unbroken cells, nuclei, and beads. The supernatant was collected and centrifuged at 100,000 × *g* for 15 min to isolate membrane fractions, which were then washed and resuspended in SEM buffer (10 mM MOPS-KOH, pH 7.2, 250 mM sucrose, 1 mM EDTA).

To purify biotinylated proteins, membrane fractions (2 mg of protein) were solubilized in 250 µl of RIPA buffer (50 mM Tris-HCl, pH 7.5, 150 mM NaCl, 1% (v/v) Triton X-100, 0.5% (w/v) sodium deoxycholate, 0.1% (w/v) SDS, 2 mM PMSF) on ice for 20 min. After centrifugation at 12,000 × *g* for 10 min, 200 µl of the supernatant was transferred to 2-ml tubes containing 1.8 ml of RIPA buffer and 20 µl of Pierce streptavidin magnetic beads (Thermo Fisher Scientific). For LC-MS/MS analysis, this procedure was scaled up fourfold using 8 mg of membrane protein. The mixtures were incubated at 4°C for 4 hours with gentle rotation. The beads were collected with a magnetic rack and sequentially washed once with RIPA buffer, 1 M KCl, 0.1 M Na₂CO₃, and urea buffer (20 mM HEPES-KOH, pH 7.5, 2 M urea), followed by two additional washes with RIPA buffer and three washes with D-PBS (-) (Fujifilm Wako Pure Chemical). For immunoblotting, bound proteins were eluted with 3× SDS sample buffer (0.375 M Tris-HCl, pH 6.8, 6.3% (w/v) SDS, 30% (w/v) sucrose, 0.01% (w/v) bromophenol blue) supplemented with 2 mM biotin. For LC-MS/MS, the beads were stored in methanol until further processing. After removing the methanol, proteins on the beads were digested in 200 µl of trypsin digestion buffer (1% (w/v) sodium deoxycholate, 1 M Urea (Thermo Fisher Scientific), 50 mM NH₄HCO₃, and 0.25 μg of sequencing-grade modified trypsin (Promega)) at 37°C for 24 h. Following digestion, 40 µl of 5% (v/v) formic acid was added, and the samples were vortexed and centrifuged at 20,000 × *g* for 10 min at room temperature. The resulting supernatant (200 µl) was extracted with 200 µl of ethyl acetate by vigorous vortexing, followed by centrifugation at 20,000 × *g* for 5 min. The upper organic phase was removed, and the aqueous phase was dried using a vacuum concentrator (Eppendorf) at 40°C. Dried peptides were dissolved in 20 µl of 0.1% (v/v) formic acid. After desalting with a C-Tip (nikkyo-tec: NTCR-KT200-C18), the samples were analyzed by LC-MS/MS using an EASY-nLC 1000 system (Thermo Fisher Scientific) coupled to a Q Exactive mass spectrometer (Thermo Fisher Scientific). Full-scan spectra were acquired in the m/z range of 380–1500 in data-dependent acquisition mode. Raw data were searched against the Swiss-Prot Saccharomyces cerevisiae protein database using Proteome Discoverer software (version 1.4, Thermo Scientific) with Mascot (version 2.8, Matrix Science, Tokyo, Japan) as the search engine. Mass tolerances for precursor and fragment ions were set to 10 ppm and 0.8 Da, respectively. Trypsin was specified as the digestion enzyme with up to one missed cleavage allowed. Methionine oxidation was set as a variable modification. The search results were filtered using Percolator with a false discovery rate (FDR) threshold of 1%. The LC-MS/MS data are summarized in Data S2.

### Fluorescence microscopy

Yeast cells in the logarithmic growth phase, cultured in YPD or SCD medium, were observed using a model IX83 microscope (Olympus) equipped with a CSU-X1 confocal unit (Yokogawa), a 100× and 1.4 numerical aperture objective lens (UPlanSApo; Olympus), and an Evolve 512 EM-CCD camera (Photometrics). LipiBlue, GFP, or mCherry were excited using a 405-nm, 488-nm or 561-nm laser (OBIS; Coherent), respectively. The confocal fluorescent sections were collected every 0.2 µm from the upper to the bottom surface of yeast cells. Image J software (NIH) was used to create maximum projection images.

For lipid droplet staining, LipiBlue (0.1 mM stock solution, Dojindo) was added to the cell culture at a final concentration of 75 nM, and the cells were incubated for 30 min. Cells were then washed twice with Milli-Q water prior to imaging.

### Phosphatase treatment

Membrane fractions were resuspended in assay buffer (300 mM sucrose, 50 mM HEPES, pH 7.5, 100 mM NaCl, 2 mM DTT, and 0.01% Brij 35) to a final protein concentration of 2 mg/ml. The suspension was treated with 200 units of Lambda Protein Phosphatase (NEB, P0753S) at 30°C for 15 min to dephosphorylate 100 µg of membrane protein. The reaction was terminated by adding an equal volume of 20% (v/v) TCA, followed by incubation on ice for 20 min and centrifugation at 20,000 × *g* for 10 min. The resulting pellet was washed with ice-cold acetone, resuspended in SDS sample buffer, and subjected to Phos-tag or SDS-PAGE.

### Immunoblotting

For immunoblotting, whole-cell extracts were prepared as follows. Yeast cells in logarithmic phase or after GS treatment were harvested at 2 OD₆₀₀ units and resuspended in 300 µl of 10% (v/v) trichloroacetic acid (TCA), followed by incubation on ice for 10 min. The cells were then centrifuged at 13,200 × *g* for 5 min at 4°C. The pellet was resuspended in 100 µl of 10% (v/v) TCA along with 100 µl of glass beads (φ0.35–0.50 mm), and disrupted by vortexing (30 seconds on, 60 seconds off, 8 cycles). After disruption, 900 µl of 10% (v/v) TCA was added to the sample and vortexed, and 800 µl of the supernatant was transferred to a new tube. The supernatant was centrifuged at 13,200 × *g* for 5 min at 4°C. The resulting pellet was resuspended in 800 µl of ice-cold acetone, followed by centrifugation at 20,000 × *g* for 5 min at 4°C. The final pellet was resuspended in 160 µl of SDS sample buffer, incubated at 37°C for 15 min, and used for SDS-PAGE. For SDS-PAGE, either homemade gels or precast gels (SuperSep™ Ace, 10-20%, Wako) were used, while Phos-tag SDS-PAGE was performed using Phos-tag precast gels (SuperSep™ Phos-tag™ (50μmol/L), 10%, Wako). After SDS-PAGE, proteins were transferred to polyvinylidene fluoride Immobilon-FL or Immobilon-P membranes (Millipore). For streptavidin blotting, PVDF membranes were blocked for 6 hours in 3% BSA blocking buffer (10 mM Tris-HCl, pH 7.5, 150 mM NaCl, 0.05% (v/v) Tween 20, 3% (w/v) BSA; Nacalai Tesque, cat# 01281-26). For standard immunoblotting, membranes were blocked for 1 hour in 1% skim milk blocking buffer (10 mM Tris-HCl, pH 7.5, 150 mM NaCl, 0.05% (v/v) Tween 20, 1% (w/v) skim milk; Morinaga, cat# 0652842). The transferred proteins were detected by fluorophore-conjugated to secondary antibodies or streptavidin such as Cy5 AffiniPure Goat Anti-Rabbit IgG (H+L) (Jackson ImmunoResearch Laboratories), Streptavidin, (Cy5), (Thermo Fisher Scientific) or goat anti-Rabbit IgG (H+L) cross-adsorbed secondary antibody conjugated to HRP (Thermo Fisher Scientific) and analyzed with Amersham Typhoon scanner (Cytiva) or Chemi Doc Touch (Bio-Rad).

### Lipid analysis

Metabolic labeling of the mevalonate pathway and analysis of the resulting radiolabeled lipids were performed essentially as described previously (*63*). Briefly, 1 µl of a saturated yeast culture was inoculated into 5 ml of YPD medium and cultivated at 30°C until the culture reached an OD₆₀₀ of 1.5. 7 OD units of cells were harvested by centrifugation and washed twice with YP medium containing 0.01% (w/v) glucose. The cells were then resuspended in 5 ml of the same medium supplemented with 1 μCi/ml of [1-¹⁴C] acetic acid, sodium salt and incubated at 30°C for 24 h.

To analyze sterol lipids, 3.5 OD units of cells were collected into 2-ml safe-lock tubes (Eppendorf) and centrifuged to pellet the cells. The pellet was resuspended in 200 µl of methanol and vortexed vigorously for 5 min at room temperature. Subsequently, 200 µl of glass beads (φ0.35–0.50 mm) were added, and the mixture was vortexed again for 5 min. After the addition of 900 µl of methanol, the samples were rotated gently for 15 min at room temperature. The samples were centrifuged at 13,200 × *g* for 5 min, and 900 µl of the supernatant was transferred to a new 2-ml tube. For saponification, 600 µl of 10% (w/v) KOH was added to the methanolic extract, and the sample was rotated at room temperature for 24 hours. After saponification, 400 µl of petroleum ether was added, and the sample was vortexed for 5 min. The mixture was centrifuged at 13,200 × *g* for 5 min, and 300 µl of the upper (organic) phase was collected into a new 2-ml tube. A second extraction was performed by adding another 400 µl of petroleum ether to the remaining aqueous phase, followed by vortexing and centrifugation under the same conditions. Then, 350 µl of the upper layer was collected and combined with the first extract. The combined organic phase was dried completely under a stream of nitrogen gas at 60°C. The dried lipids were dissolved in 30 µl of petroleum ether. Twenty microliters of the sample were spotted onto a TLC plate (MACHEREY-NAGEL, #810123). The plate was developed in solvent A (benzene:ethyl acetate = 100:20) to a height of ∼10 cm from the bottom, air-dried for 10 min, and developed again in the same solvent to the same distance. After drying, the plate was further developed in solvent B (petroleum ether:diethyl ether:acetic acid = 95:5:1) to ∼15 cm and dried. Radiolabeled lipids were detected by autoradiography using an Amersham Typhoon scanner (Cytiva).

To analyze phospholipids, 3.0 OD_600_ units of cells were collected into 2-ml safe-lock tubes (Eppendorf) and centrifuged to pellet the cells. The pellet was resuspended in 300 µl of methanol and vortexed vigorously for 5 min at room temperature. Subsequently, 200 µl of glass beads (φ0.35–0.50 mm) were added, and the mixture was vortexed again for 20 min. After the addition of 600 µl of chloroform, the samples were vortexed for 5 min at room temperature. The samples were centrifuged at 13,200 × *g* for 5 min, and 750 µl of the supernatant was transferred to a new 2-ml tube. Then, 200 µl of 0.1M NaCl, 0.1 M HCl was added to the samples and further vortexed for 5 min at room temperature. After centrifugation at 210 × *g* for 5 min at room temperature, the upper phase was removed, and the lower phase was dried completely under a stream of nitrogen gas at 60°C. The dried lipids were dissolved in 30 µl of chloroform. Twenty microliters of the sample were spotted onto a TLC plate (MACHEREY-NAGEL, #810123), which was developed in solvent C (chloroform:ethanol:triethylamine:water = 30:35:35:5) to a height of approximately 15 cm from the bottom. Radiolabeled lipids were detected by autoradiography using an Amersham Typhoon scanner (Cytiva).

### Statistical analyses

Data are shown as mean with standard error of the mean as indicated in the figure legends. Student’s t-test were performed for the statistical analyses using Prism 10 software (GraphPad).

## Acknowledgments

We thank M. Hashimoto for her great technical assistance. We are grateful to the members of the Tamura laboratory for helpful discussion. We also thank Prof. Toshiya Endo (Kyoto Sangyo University) for providing antibodies and plasmids.

## Author contributions

Conceptualization: SF, YT

Methodology: SF, YT

Investigation: SF, YT

Visualization: SF, YT

Supervision: YT

Writing—original draft: YT

Writing—review & editing: SF, YT

## Data and materials availability

All data are available in the main text or the supplementary materials. Requests for resources should be directed to and will be fulfilled by the corresponding author, Yasushi Tamura.

---

**Fig. S1.**
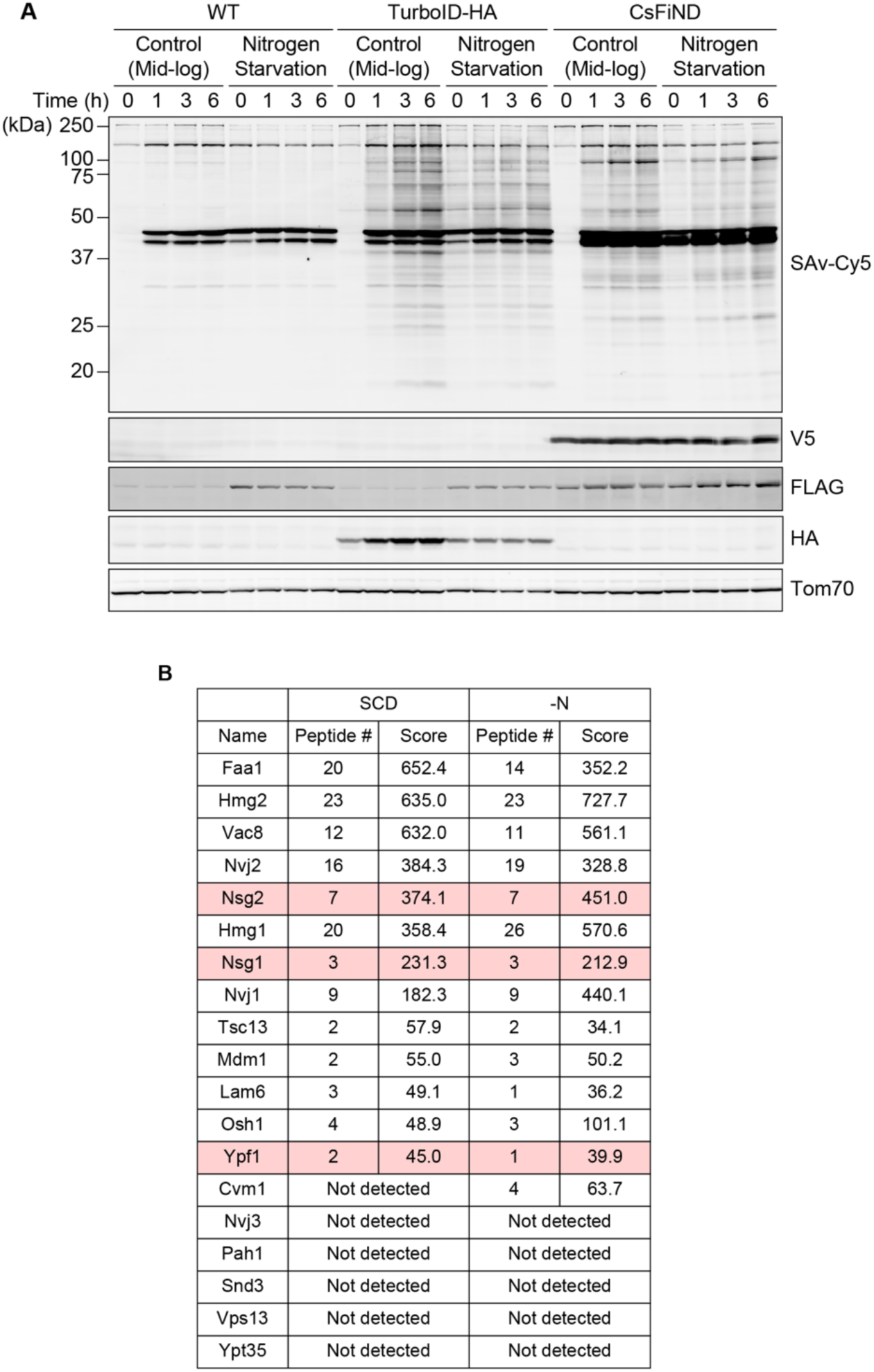
Identification of NVJ-localized proteins using CsFiND. (**A**) Wild-type yeast cells and yeast cells expressing CsFiND proteins or cytosolic TurboID-HA were cultured in SCD medium until logarithmic phase, and then split into two halves. One is further cultivated in SCD medium containing biotin. Another half is shifted to SD-N medium containing biotin at a final concentration of 50 μM for the indicated hours. Total cell lysates were analyzed by streptavidin blotting or immunoblotting using the indicated antibodies. (**B**) Summary table showing protein scores obtained from MASCOT analysis of LC-MS/MS data for known NVJ proteins, as well as Ypf1, Nsg1, and Nsg2.

**Fig. S2.**
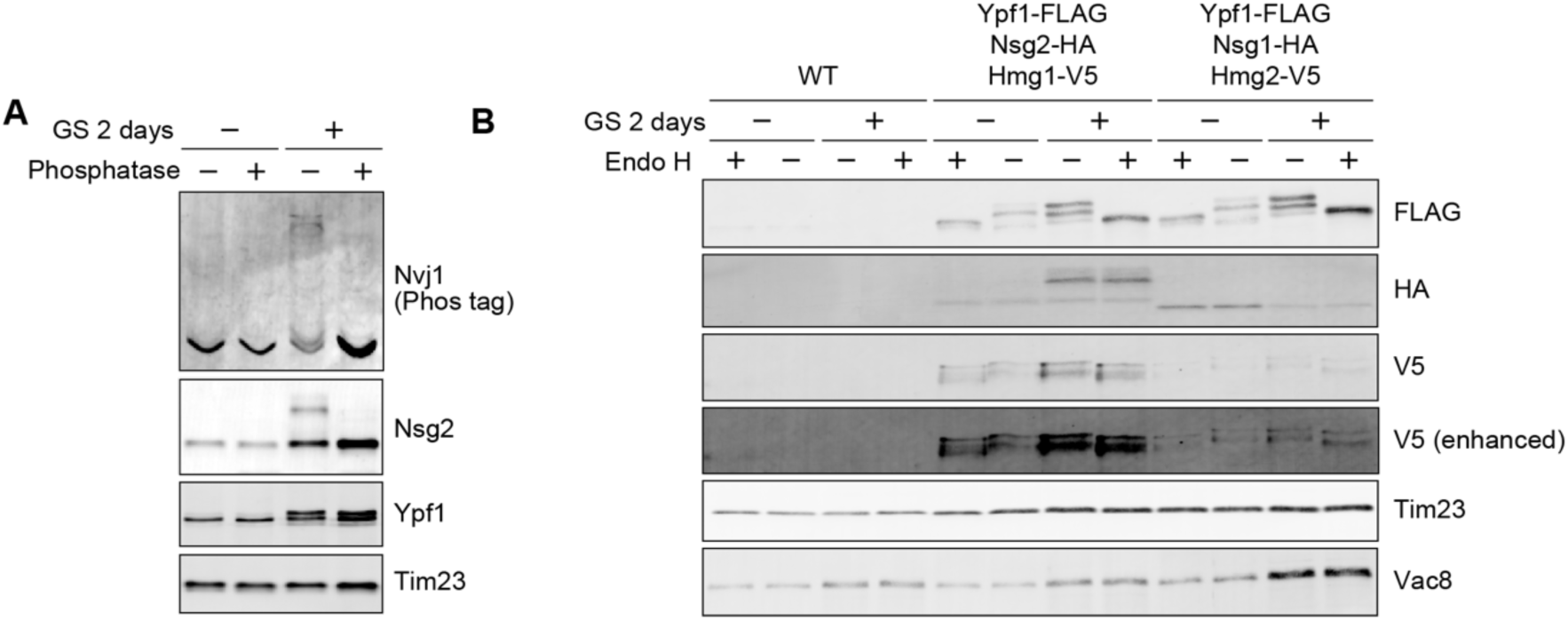
Nvj1 and Nsg2 are phosphorated in a GS-dependent manner. (**A**) Membrane fractions isolated from yeast cells with or without 2 days of GS treatment were treated with λ protein phosphatase. The samples were analyzed by Phos-tag or SDS-PAGE followed by immunoblotting. (**B**) Membrane fractions isolated from the indicated yeast cells cultured with or without 2 days of GS, and those marked as Endo H+ were subjected to Endo H digestion before immunoblot analysis.

**Fig. S3.**
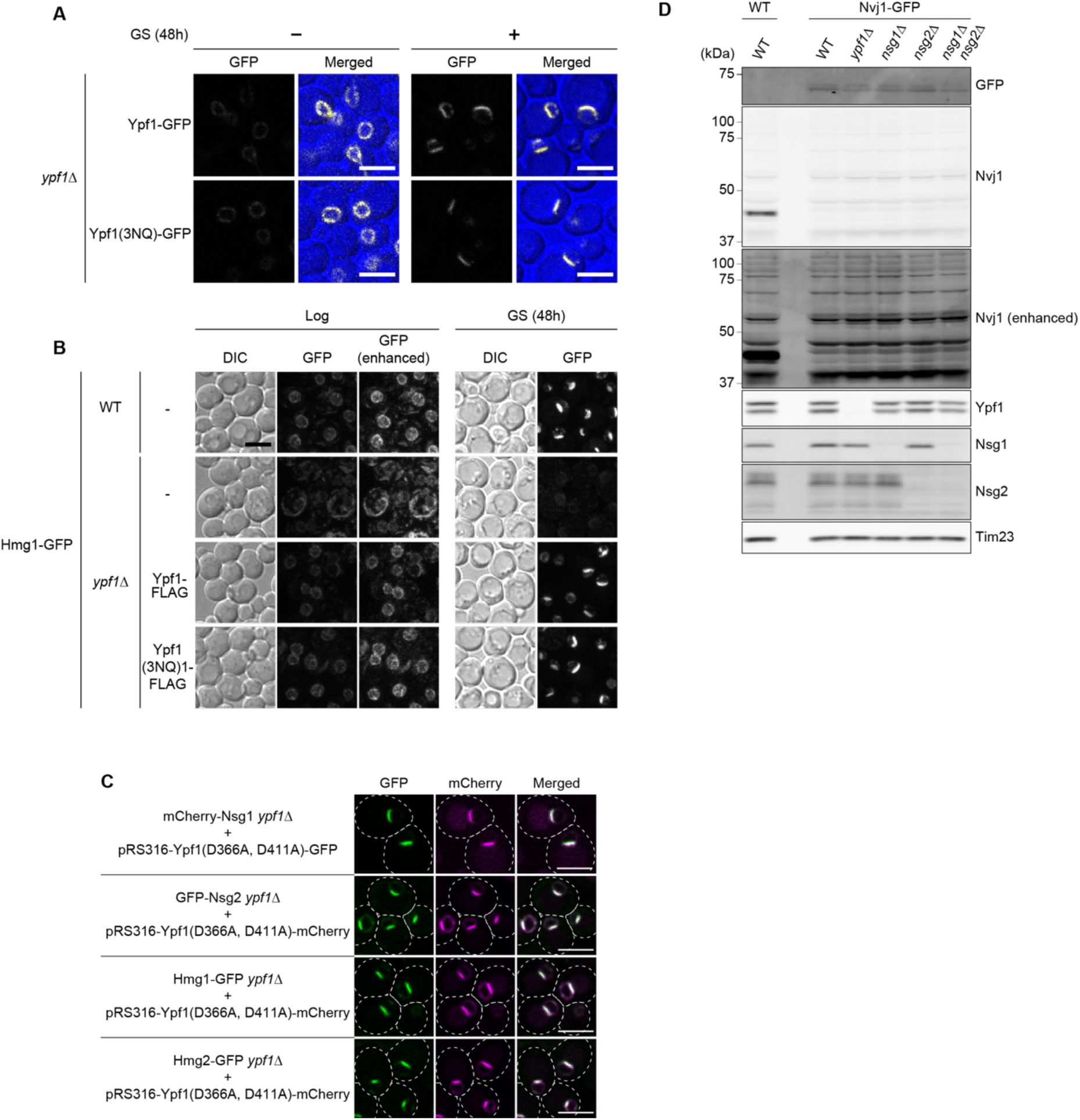
Neither the glycosylation of Ypf1 nor its peptidase activity is required for GS– dependent remodeling of the NVJ. (**A**) *ypf1*Δ cells harboring a *CEN* plasmid expressing either Ypf1-GFP or the glycosylation-deficient mutant Ypf1-3NQ-GFP were observed by confocal fluorescence microscopy after 2 days of GS. (**B**) Wild-type and *ypf1*Δ cells expressing Hmg1-GFP and harboring either an empty vector (–), Ypf1-FLAG, or Ypf1-3NQ-FLAG-expressing plasmid were observed by confocal fluorescence microscopy after 2 days of GS. (**C**) A *CEN* plasmid encoding catalytically inactive Ypf1-D366A/D411A-mCherry was introduced into *ypf1*Δ cells expressing mCherry-Nsg1, GFP-Nsg2, Hmg1-GFP, or Hmg2-GFP, and cells were analyzed by confocal fluorescence microscopy after 1 day of GS. (**D**) Total cell lysates from the indicated strains after 1 day of GS were analyzed by immunoblotting. All images in (**A**)-(**C**) are single focal plane. Scale bar, 5 μm.

**Figure S4.**
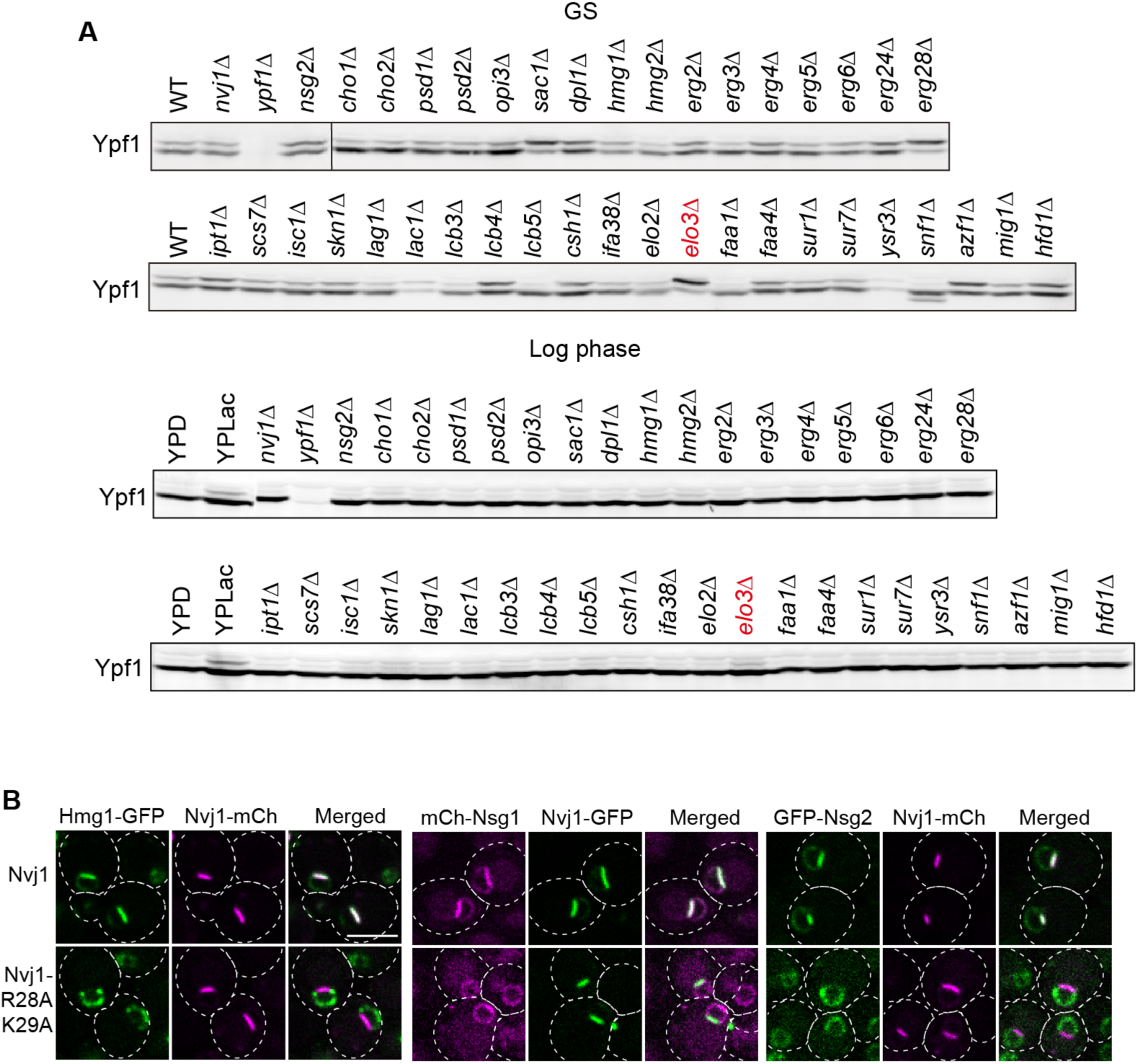
Deletion of ELO3 promotes GS–dependent glycosylation of Ypf1. (**A**) Glycosylation patterns of Ypf1 were analyzed by immunoblotting using total cell lysates prepared from the knockout strains listed in Table S5. Lysates were obtained from cells harvested either during logarithmic growth (Log) or after 1 day of GS. (**B**) NVJ accumulation of Hmg1, Nsg1, and Nsg2 in cells expressing wild-type or mutant Nvj1 was observed by confocal fluorescence microscopy after 1 day of GS. All images are single focal plane. Scale bar, 5 μm.

**Fig. S5.**
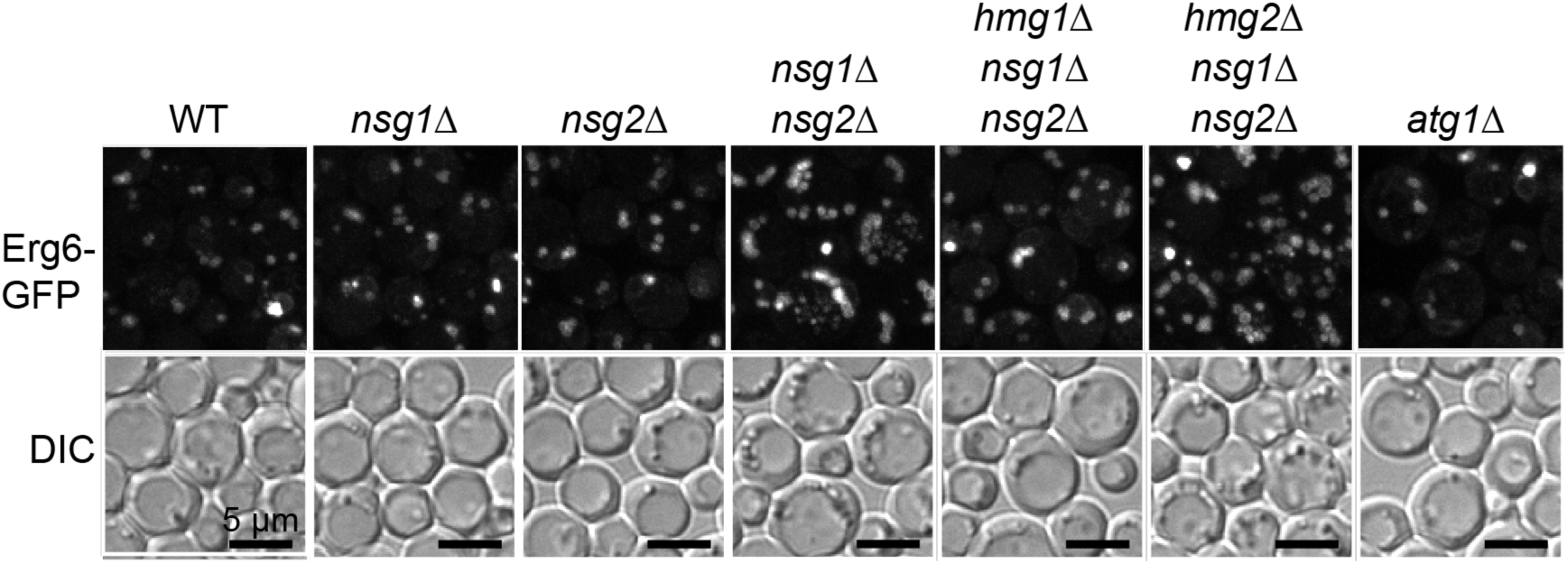
LDs accumulate in the absence of Nsg1 and Nsg2. Wild-type and the indicated cells expressing Erg6-GFP were observed by confocal fluorescence microscopy after 4 days of GS. Maximum projection images reconstituted from z-stack are shown. Scale bars, 5 μm.

**Table S1.**
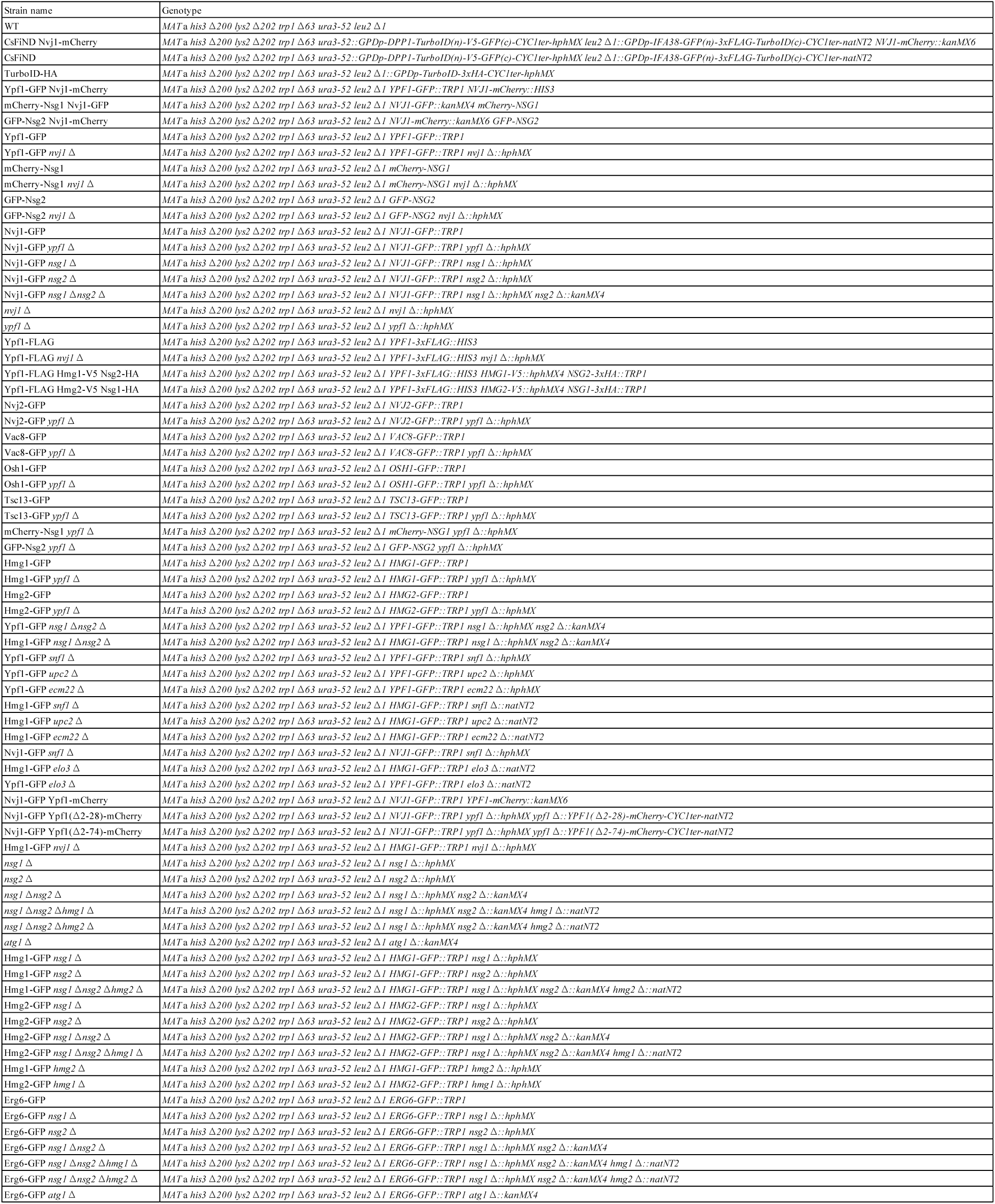
Yeast strains used in this study.

**Table S2.** Yeast gene knock out strains used in this study.

| Strain name | Genotype |
| --- | --- |
| WT | *MATa his3 Δ1 leu2 Δ0 met15 Δ0 ura3 Δ0* |
| *nvj1 Δ* | *MATa his3 Δ1 leu2 Δ0 met15 Δ0 ura3 Δ0 nvj1 Δ::kanMX4* |
| *ypf1 Δ* | *MATa his3 Δ1 leu2 Δ0 met15 Δ0 ura3 Δ0 ypf1 Δ::kanMX4* |
| *nsg2 Δ* | *MATa his3 Δ1 leu2 Δ0 met15 Δ0 ura3 Δ0 nsg2 Δ::kanMX4* |
| *cho1 Δ* | *MATa his3 Δ1 leu2 Δ0 met15 Δ0 ura3 Δ0 cho1 Δ::kan<sup>r</sup>* |
| *cho2 Δ* | *MATa his3 Δ1 leu2 Δ0 met15 Δ0 ura3 Δ0 cho2 Δ::kan<sup>r</sup>* |
| *psd1 Δ* | *MATa his3 Δ1 leu2 Δ0 met15 Δ0 ura3 Δ0 psd1 Δ::kanMX4* |
| *psd2 Δ* | *MATa his3 Δ1 leu2 Δ0 met15 Δ0 ura3 Δ0 psd2 Δ::kanMX4* |
| *opi3 Δ* | *MATa his3 Δ1 leu2 Δ0 met15 Δ0 ura3 Δ0 opi3 Δ::kan<sup>r</sup>* |
| *sac1 Δ* | *MATa his3 Δ1 leu2 Δ0 met15 Δ0 ura3 Δ0 sac1 Δ::kanMX4* |
| *dpl1 Δ* | *MATa his3 Δ1 leu2 Δ0 met15 Δ0 ura3 Δ0 dpl1 Δ::kanMX4* |
| *hmg1 Δ* | *MATa his3 Δ1 leu2 Δ0 met15 Δ0 ura3 Δ0 hmg1 Δ::kanMX4* |
| *hmg2 Δ* | *MATa his3 Δ1 leu2 Δ0 met15 Δ0 ura3 Δ0 hmg2 Δ::kanMX4* |
| *erg2 Δ* | *MATa his3 Δ1 leu2 Δ0 met15 Δ0 ura3 Δ0 erg2 Δ::kanMX4* |
| *erg3 Δ* | *MATa his3 Δ1 leu2 Δ0 met15 Δ0 ura3 Δ0 erg3 Δ::kanMX4* |
| *erg4 Δ* | *MATa his3 Δ1 leu2 Δ0 met15 Δ0 ura3 Δ0 erg4 Δ::kanMX4* |
| *erg5 Δ* | *MATa his3 Δ1 leu2 Δ0 met15 Δ0 ura3 Δ0 erg5 Δ::kanMX4* |
| *erg6 Δ* | *MATa his3 Δ1 leu2 Δ0 met15 Δ0 ura3 Δ0 erg6 Δ::kanMX4* |
| *erg24 Δ* | *MATa his3 Δ1 leu2 Δ0 met15 Δ0 ura3 Δ0 erg24 Δ::kanMX4* |
| *erg28 Δ* | *MATa his3 Δ1 leu2 Δ0 met15 Δ0 ura3 Δ0 erg28 Δ::kanMX4* |
| *ipt1 Δ* | *MATa his3 Δ1 leu2 Δ0 met15 Δ0 ura3 Δ0 ipt1 Δ::kanMX4* |
| *scs7 Δ* | *MATa his3 Δ1 leu2 Δ0 met15 Δ0 ura3 Δ0 scs7 Δ::kanMX4* |
| *isc1 Δ* | *MATa his3 Δ1 leu2 Δ0 met15 Δ0 ura3 Δ0 isc1 Δ::kanMX4* |
| *skn1 Δ* | *MATa his3 Δ1 leu2 Δ0 met15 Δ0 ura3 Δ0 skn1 Δ::kanMX4* |
| *lag1 Δ* | *MATa his3 Δ1 leu2 Δ0 met15 Δ0 ura3 Δ0 lag1 Δ::kanMX4* |
| *lac1 Δ* | *MATa his3 Δ1 leu2 Δ0 met15 Δ0 ura3 Δ0 lac1 Δ::kanMX4* |
| *lcb3 Δ* | *MATa his3 Δ1 leu2 Δ0 met15 Δ0 ura3 Δ0 lcb3 Δ::kanMX4* |
| *lcb4 Δ* | *MATa his3 Δ1 leu2 Δ0 met15 Δ0 ura3 Δ0 lcb4 Δ::kanMX4* |
| *lcb5 Δ* | *MATa his3 Δ1 leu2 Δ0 met15 Δ0 ura3 Δ0 lcb5 Δ::kanMX4* |
| *csh1 Δ* | *MATa his3 Δ1 leu2 Δ0 met15 Δ0 ura3 Δ0 csh1 Δ::kanMX4* |
| *ifa38 Δ* | *MATa his3 Δ1 leu2 Δ0 met15 Δ0 ura3 Δ0 ifa38 Δ::kanMX4* |
| *elo2 Δ* | *MATa his3 Δ1 leu2 Δ0 met15 Δ0 ura3 Δ0 elo2 Δ::kanMX4* |
| *elo3 Δ* | *MATa his3 Δ1 leu2 Δ0 met15 Δ0 ura3 Δ0 elo3 Δ::kanMX4* |
| *faa1 Δ* | *MATa his3 Δ1 leu2 Δ0 met15 Δ0 ura3 Δ0 faa1 Δ::kanMX4* |
| *faa4 Δ* | *MATa his3 Δ1 leu2 Δ0 met15 Δ0 ura3 Δ0 faa4 Δ::kanMX4* |
| *sur1 Δ* | *MATa his3 Δ1 leu2 Δ0 met15 Δ0 ura3 Δ0 sur1 Δ::kanMX4* |
| *sur7 Δ* | *MATa his3 Δ1 leu2 Δ0 met15 Δ0 ura3 Δ0 sur7 Δ::kanMX4* |
| *ysr3 Δ* | *MATa his3 Δ1 leu2 Δ0 met15 Δ0 ura3 Δ0 ysr3 Δ::kanMX4* |
| *snf1 Δ* | *MATa his3 Δ1 leu2 Δ0 met15 Δ0 ura3 Δ0 snf1 Δ::kanMX4* |
| *azf1 Δ* | *MATa his3 Δ1 leu2 Δ0 met15 Δ0 ura3 Δ0 azf1 Δ::kanMX4* |
| *mig1 Δ* | *MATa his3 Δ1 leu2 Δ0 met15 Δ0 ura3 Δ0 mig1 Δ::kanMX4* |
| *hfd1 Δ* | *MATa his3 Δ1 leu2 Δ0 met15 Δ0 ura3 Δ0 hfd1 Δ::kanMX4* |

**Table S3.**
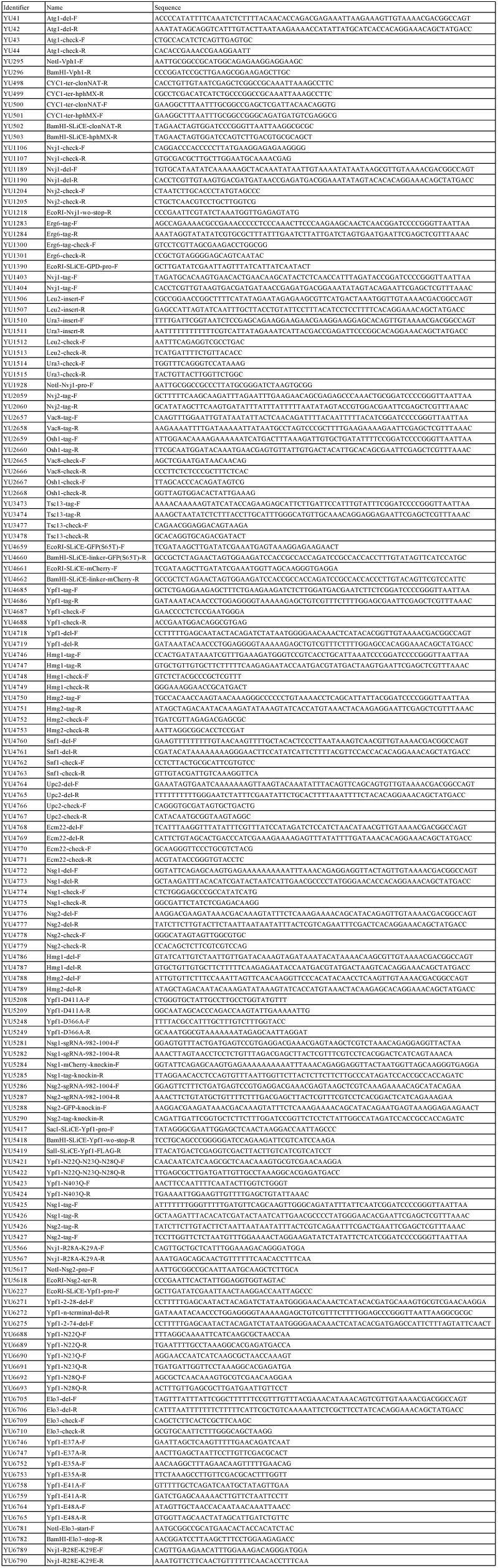
Yeast gene knock out strains used in this study.

